# Automated sample preparation with SP3 for low-input clinical proteomics

**DOI:** 10.1101/703413

**Authors:** Torsten Müller, Mathias Kalxdorf, Rémi Longuespée, Daniel N. Kazdal, Albrecht Stenzinger, Jeroen Krijgsveld

## Abstract

High-throughput and streamlined workflows are essential in clinical proteomics for standardized processing of samples originating from a variety of sources, including fresh frozen tissue, FFPE tissue, or blood. To reach this goal, we have implemented single-pot solid-phase-enhanced sample preparation (SP3) on a liquid handling robot for automated processing (autoSP3) of tissue lysates in a 96-well format, performing unbiased protein purification and digestion, and delivering peptides that can be directly analyzed by LCMS. AutoSP3 eliminates hands-on time and minimizes the risk of error, reduces variability in protein quantification and improves longitudinal performance and reproducibility. We demonstrate the distinguishing ability of autoSP3 to process low-input samples, reproducibly quantifying 500-1000 proteins from 100-1000 cells (<100 ng protein). Furthermore, we applied this approach to a cohort of clinical FFPE pulmonary adenocarcinoma (ADC) samples, and recapitulate their separation into known histological growth patterns based on proteome profiles. Collectively, autoSP3 provides a generic, scalable, and cost-effective pipeline for routine and standardized proteomic sample processing that should enable reproducible proteomics in a broad range of clinical and non-clinical applications.

## Introduction

Mass spectrometry (MS)-based proteomic technologies have matured to allow robust, reliable, and comprehensive proteome profiling across thousands of proteins in cells and tissues. This is the result of parallel developments in mass spectrometric instrumentation that continues to gain speed and sensitivity, in liquid chromatographic technology to separate peptides directly interfaced with MS, and in data analysis pipelines for reliable protein identification and quantification. In addition, various workflows have been developed for comparative analyses across many samples, e.g. using isobaric labels allowing sample multiplexing, or using label-free approaches and short liquid chromatography (LC) gradients. Collectively this has propelled proteomic studies in multiple areas of basic and mechanistic biology, using deep and quantitative proteomic profiles to understand spatial and temporal aspects of proteome organization and dynamics in a wide variety of static or perturbed conditions^1^. In addition, the speed, sensitivity, robustness, and general accessibility of present-day proteomic technologies have an increasing appeal for clinical applications, for various reasons: i) underlying mechanisms of many diseases are still unclear, where proteome-level information will increase mechanistic insight of (patho)physiological processes; ii) proteins are the primary targets of almost all current drugs, and insight in their function will help to understand how drugs impact on cellular processes; iii) for many diseases there is a persistent lack of powerful protein biomarkers for diagnostic, prognostic, or predictive purposes.

Still, successful implementation of proteomics in a clinical environment has not materialized yet, primarily because of additional requirements that need to be met on top of those in a research environment alluded to above (e.g. proteome coverage, sensitivity). This mostly pertains to i) the ability to analyze many (possibly hundreds) of samples in an un-interrupted fashion in order to achieve sufficient statistical power across patients, ii) simplify the workflow, thereby removing the need for personnel with specific technical skills in proteomics, iii) achieving an acceptable turnaround time from receiving samples to the generation of a complete proteome profile analysis, and iv) cost-effectiveness of the workflow. Most of these bottlenecks can be resolved simultaneously by automation, avoiding manual handling and thereby eliminating the risk of error and variability, while at the same time enabling longitudinal standardization irrespective of the number of samples. Although LCMS has nowadays been sufficiently standardized to achieve excellent performance across hundreds of samples^2^, preceding sample preparation is still highly cumbersome, involving multiple steps to extract, purify, and digest proteins before subsequent LCMS. In an ideal scenario, this procedure should be streamlined into an automated pipeline that accepts processing conditions for any sample type, thereby facilitating universal applicability. Despite the range of existing sample preparation methods^3–10^, very few satisfy these demands to universally accommodate the different requirements imposed by various clinical tissue types, e.g. blood cells can be lysed under more gentle conditions than fresh frozen tissue, while formalin-fixed and paraffin-embedded (FFPE) tissue requires harsh detergent-based methods to efficiently extract proteins. Among the most popular sample preparation methods, stage tips^6^, and its derivative iST^8^, do not tolerate detergents thereby restricting their generic use. In addition, most methods involve extensive manual handling and procedures such as filtration^3, 8, 10^, centrifugation^3, 8, 10^, precipitation^5^, and electrophoresis^4^ that are difficult to standardize and unsuitable for automation.

We recently introduced single-pot solid-phase-enhanced sample preparation (SP3) (Hughes et al., MSB 2014; Hughes et al., Nat Protocols 2019) for unbiased protein retrieval and purification^11, 12^. The method utilizes paramagnetic beads in the presence of an organic solvent (>50% ACN or ethanol) to promote protein binding to the beads, allowing extensive washing to eliminate contaminants, including detergents such as SDS and TritonX. After release of proteins off the beads in an aqueous buffer, proteolytic digestion produces clean peptides that can be directly injected for analysis by LCMS. Another distinguishing feature of SP3 is its efficient protein recovery, facilitating low-input applications while maintaining deep proteome coverage. The combined characteristics of scalability, tolerance to detergents, speed and ease of operation, qualify SP3 as a universal methodology that has enabled a wide variety of applications, including those involving ‘difficult’ sample types as diverse as FFPE tissue^13, 14^ and historical bones^15^. In addition, SP3 especially performs well for low-input applications^16^ e.g. allowing an analysis of single human oocytes^17^, and micro-dissected tissue^18, 19^.

A property of SP3 that has not been fully exploited yet is the paramagnetic nature of the beads, which allow automation of the procedure on a robotic liquid handling platform. In genomics, automated sample preparation using magnetic beads has been introduced already several years ago^20^, and is now commonly used for library generation through kits available form many vendors. Automated sample preparation is far less common in proteomics, and is restricted to cases where detergents can be avoided (iST, plasma proteomics^21^), or for the enrichment of specific sub-proteomes (e.g. AssayMap to purify phosphorylated peptides^22^) or for protein digestion and peptide clean-up^23^.

In this study, we implemented SP3 on a liquid handling system, in order to build a generic, automated, and scalable 96-well format proteomics pipeline that performs all handling steps starting from a tissue lysate and delivering protein digests for MS analysis. In addition, we verified performance stability over a period of several weeks and demonstrated the ability to reproducibly handle low-input samples, down to low ng samples containing 100 cells or less. To demonstrate the integration of automated SP3 in a realistic clinical scenario, we analyzed a cohort of pulmonary adenocarcinoma (ADC) samples, successfully associating pathological tumor growth patterns that have strong prognostic implications^24^ with distinct proteomic signatures. Collectively, automated SP3 addresses an unmet need in streamlining and hands-free sample processing from tissues to peptides. This provides an attractive and cost-effective solution for routine, comprehensive clinical studies, easing the introduction of translational proteomic research with minimal hands-on time and low sample consumption.

## Experimental Methods

### Chemicals

HeLa cells were purchased from ATCC (Wesel, Germany). Trypsin-EDTA (0.25%), 100 × glutamine stock solution, Penicillin-Streptomycin (P&S) mix, Fetal Bovine serum (FBS), and Dulbecco’s modified Eagle Medium (DMEM) with high glucose and no glutamine were obtained from Life Technologies (Darmstadt, Germany). Protease inhibitor cocktail (PIC), Tris(2-carboxyethyl)phosphine (TCEP), and chloroacetamide (CAA) were obtained from Sigma-Aldrich (Steinheim, Germany). Benzonase and ethanol (EtOH) were from Merck (Darmstadt, Germany). Acetonitrile (ACN) was obtained from Fisher Scientific (Schwerte, Germany). All solvents were MS grade. Sodium-dodecylsulfate (SDS) was from Applichem (Darmstadt, Germany) and phosphate buffered saline (PBS) was from Biowest (Darmstadt, Germany). Ammonium bicarbonate (ABC) was purchased from Fluka Analytical (Munich, Germany). LCMS-grade water, Trifluoroacetic acid (TFA), and formic acid (FA) were obtained from Biosolve Chemicals (Dieuze, France). Sequencing grade modified trypsin with acetic acid resuspension buffer was obtained from Promega (Madison, WI, USA). Paramagnetic beads for SP3 (Sera-Mag Speed Beads A and B) were purchased from Fisher Scientific (Schwerte, Germany).

### Cell Culture of HeLa Cells

HeLa cells were cultured in regular DMEM medium supplemented with 10% fetal bovine serum, 1% of a 100 × penicillin and streptomycin mix, and 1% of 100 × glutamine stock solution (Gibco). Upon establishment of a stable culture cells were harvested using trypsin and counted using Bio-Rad TC20 automated cell counter. Cell pellets were stored at -80 °C until further use.

For showing the use of the Bravo application starting from limited, small numbers of cells, HeLa cells were harvested, counted, resuspended in lysis buffer (1% SDS, 100 mM ABC pH 8.5), and directly transferred to a 96-well plate. The total volume for different numbers of cells was adjusted using lysis buffer (1% SDS, 100 mM ABC pH 8.5). The entire 96-well plate was sonicated in a waterbath for 10 minutes, followed by Benzonase (∼40 Units) enzymatic cleavage of DNA and RNA for 15 minutes at 37°C. Subsequently, the buffer was adapted to a final concentration of 1% SDS, 100 mM ABC, 10 mM TCEP, and 40 mM CAA including protease inhibitor cocktail (PIC)) before incubation for 5 minutes at 95°C. The plate was allowed to cool to room temperature before it was transferred to the Bravo deck for the SP3 processing as described in the “automated SP3 protocol*”* section.

### HeLa Protein Standard Preparation

Cell pellets of ∼11.9 million cells were resuspended in 1 mL of lysis buffer (1% SDS, 100 mM ABC pH 8.5, and 50 μL 25x PIC) and probe sonicated for 5 times 20 seconds at a frequency of 10% using a Branson Sonifier. Cell lysates were kept on ice in-between cycles to avoid overheating. DNA or RNA contaminants were cleaved using 250 Units of Benzonase for 15 minutes at 37°C and 750 rpm. Subsequently, the buffer was adapted to a final concentration of 1% SDS, 100 mM ABC, 10 mM TCEP, and 40 mM CAA including protease inhibitor cocktail (PIC)) before incubation for 5 minutes at 95°C CHB-T2-D ThermoQ, Hangzhou BIOER Technologies). Reduced and alkylated proteins were quantified using a BCA assay and stored at -20°C until further use in manual and automated SP3 processing.

### Pulmonary Adenocarcinoma Sample Collection

All pulmonary adenocarcinoma (ADC) specimens used for this study were obtained from the Thoraxklinik at Heidelberg University and diagnosed according to the criteria of the 2015 WHO Classification of lung tumors at the Institute of Pathology at Heidelberg University^25^. Tissue procession to formalin-fixed and paraffin-embedded (FFPE) tissue sections was carried out by the tissue bank of the National Center for Tumor Diseases (NCT; project: # 1746; # 2818) in accordance with its ethical regulations approved by the local ethics committee.

A multiregional sample set consisting of 2-4 samples of eight tumors was constructed as described previously^26^. In short, a formalin fixed central section of each tumor was segmented in into multiple 5 × 5 mm regions according to a Cartesian grid. Ink marks ensured the retention of the original orientation of each segment during sample processing. Tumor regions considered for analysis were selected in accordance with the tumor size (the larger the tumor the more regions), different histological growth patterns as well as sufficient tumor cell content (≥ 10%). The histological growth pattern with predominant portion in each segment was determined by an experienced pathologist. For each tumor, two to four different growth patterns were excised. Samples were analyzed in replicates using one 5 µm section after deparaffinization as input, respectively. For deparaffinization, the sections were incubated for 20 minutes at 80°C followed by three times 8 minutes incubation in Xylol and EtOH, consecutively. Finally, the sections were incubated in ddH_2_O for 30 minutes before the tissue was scratched off and collected in a well. Replicates were excised as consecutive cuts of the same region having the highest possible similarity.

### SP3 Protocol

As a reference, the SP3 protocol was carried out manually as described before (Hughes 2019)^12^. In brief, 10 μg of extracted HeLa protein were added to PCR tubes in a total volume of 10 μL lysis buffer (1% SDS, 100 mM ABC pH 8.5). Magnetic beads were prepared by combining 20 μL of both, Sera-Mag Speed Beads A and B (Fisher Scientific, Germany), and wash them one time with 160 μL and two times with 200 μL ddH_2_O, and re-suspend them in 20 μL ddH_2_O for a final working concentration of 100 μg/μL. 2 μL of pre-washed magnetic beads as well as 12 μL 100% acetonitrile (ACN) were added to each sample to reach a final concentration of 50% ACN. Protein binding to the beads was allowed for 18 minutes, followed by 2 minutes incubation on a magnetic rack to immobilize beads. The supernatant was removed and beads were washed two times with 200 μL of 80% ethanol (EtOH) and one time with 180 μL of 100% ACN. Beads were resuspended in 15 μL of 100 mM ABC and sonicated for 5 minutes in a water bath. Finally, sequencing-grade trypsin was added in an enzyme:protein ratio of 1:20 (5 μL of 0.1 μg/μL trypsin in ddH_2_O), and beads were pushed from the tube walls into the solution to ensure efficient digestion. Upon overnight incubation at 37°C and 1000 rpm in a table-top thermomixer, samples were acidified by adding 5 μL of 5% TFA and quickly vortexed. Beads were immobilized on a magnetic rack and peptides were recovered by transferring the supernatant to new PCR tubes. Samples were acidified by adding 75 μL 0.1% FA to reach a peptide concentration of approximately 1 μg/10 μL. MS injection-ready samples were stored at -20 C.

### Automated SP3 Protocol (autoSP3)

In the automated version of the SP3 protocol, the Bravo system is programmed to process 96 samples simultaneously, carrying out all handling steps including reduction and alkylation of proteins, aliquoting of magnetic beads, protein clean-up by SP3, protein digestion, and peptide recovery. The core SP3 protocol is available in combination with reduction and alkylation either as a single-step using a TCEP/ CAA mix for 5 minutes at 95 C (**Supplementary Protocol A**) or as a two-step protocol using, for example, DTT/ CAA consecutively with 30 minutes incubation for each reaction (**Supplementary Protocol B**, Supplemental Figure 1). Both are provided as protocol files (*.vzp file) in **Supplementary Data A** (TCEP/ CAA) and **B** (DTT/ CAA) for direct use on the Bravo platform. A shortened version is available that consists of the core SP3 protocol while omitting on-deck reduction and alkylation (**Supplementary Protocol C** & *.vzp file in **Supplementary Data C**), saving time due to slow heating of the heating block (altogether taking 1 hour to heat and cool), instead performing this off-deck (taking 5 minutes and 30 seconds to reach working temperature and 5 minutes for incubation) in a PCR thermocycler (CHB-T2-D ThermoQ, Hangzhou BIOER Technologies) prior to initiation of the automated protocol. In addition, the PCR thermocycler provides lid heating, which prevents any unwanted evaporation or variation in sample volume. This latter protocol (**Supplementary Data C**) was used in the work presented here. Each protocol is designed for a starting sample volume of 10 μL, which can easily be varied in the protocol files to add respective amounts of organic solvent to reach higher than 50% and to remove the resulting volume after protein binding. Next, either protocol A, B, or C (Supplementary Figure 1) aliquot 5 μL of a suspension of washed magnetic beads to protein samples previously collected in a 96-well plate. Different to the manual protocol (bead working concentration 100 μg/μL), the suspension of washed beads is prepared to have a working concentration of 50 μg/ μL to allow more robust pipetting. Next, the respective volume of 100% ACN (20 μL in A; 25 μL in B, 15 μL in C) is added to each sample followed by 18 minutes incubation off the magnetic rack with cycles of agitation at 1500 rpm and 100 rpm for 30 seconds and 90 seconds, respectively. Upon binding of the proteins to the beads, the sample plate is incubated on the magnetic rack for further 5 minutes to allow magnetic trapping of beads inside each well. Here, the beads will form a ring at the wall of each well, slightly above the bottom. The removal of any supernatant in the protocol is performed using well-specific tips in two consecutive steps to ensure complete liquid removal. Next, beads are washed two times with 200 μL of 80% EtOH and one time with 171.5 μL of 100% ACN. Due to the limited 200 μL pipetting volume of the Bravo and the limited reagent space, the respective washing volumes of 80% EtOH and 100% ACN were added in 4 and 7 consecutive steps of 50 μL and 24.5 μL, respectively, with in-between shaking at 500 rpm or 250 rpm for 30 seconds. Upon removal of residual washing solvents, the beads are resuspended in 35 μL of 100 mM ABC and 5 μL of 0.05 μg/ μL pre-prepared trypsin in 50 mM acetic acid to avoid autolysis. Of note, in the dilution series experiments the trypsin amount was reduced to avoid abundant peptide features resulting from its autolysis. In a final shaking step at 1500 rpm for 60 seconds the trypsin solution is mixed with the sample and the plate is transferred to the heating deck position for incubation at 37°C. Subsequently, the plate was manually sealed and transferred to a PCR cycler to avoid lid condensation during a 4-hour incubation at 37°C. Next, after completion of either protocol A, B, or C and exchange of used pipette tips a short protocol is provided for peptide acidification and recovery of LCMS injection-ready samples to a new 96-well plate (**Supplementary Protocol D** & *.vzp file in **Supplementary Data D**). Alternatively, as used in this study, peptide acidification and recovery can be performed manually. Therefore, each sample was acidified by adding 5 μL of 5% TFA solution, sonicated in a water bath for 5 minutes to swirl the settled beads, and incubated on a magnetic rack for further 2 minutes. Finally, the peptide-containing supernatant was recovered into a new 96-well plate without transferring of the beads. If necessary, samples were either diluted, or directly frozen at -20°C until MS acquisition. Optionally peptide quantification assays (colorimetric assay kit, Thermo Scientific) were carried out using the Bravo liquid handling system.

### Quantitative Proteomics Analysis of FFPE Tissue

For proteomic analysis, 5 μm FFPE tissue sections were collected in stripes of 8 PCR tubes, centrifuged at 15.000 × g for 10 minutes to ensure that FFPE slices are at the bottom of the tube, and stored at 4°C until further processing. Next, each tissue section was carefully reconstituted in 20 μL lysis buffer (4% SDS, 100 mM ABC pH 8.5), sonicated at 4°C for 25 cycles of 30 seconds on and 30 seconds off in a Pico Bioruptor, and heated for 1 hour at 95°C. Samples were spun down and subjected to a second round of sonication and heating. The Pico Bioruptor was equipped with a house-made tube holder, which allows the simultaneous processing of 28 samples. Subsequently, PCR tubes were centrifuged at 15.000 × g for 3 minutes and the buffer was adjusted to a final concentration of 1% SDS, 100 mM ABC, 10 mM TCEP, and 40 mM CAA including protease inhibitor cocktail (PIC). Samples were heated for 5 minutes at 95°C to denature proteins and to reduce and alkylate cysteine residues. Cooled to RT and again centrifuged at 15.000 × g for 3 minutes, 10 μL of each sample was further processed by our automated SP3 sample clean-up procedure, as described above. Here, protein digestion was allowed for 16 hours overnight before stopping the reaction by acidification to 0.5% with TFA. The peptide-containing supernatant was recovered to a new 96-well plate without transferring of the beads. MS injection-ready samples were stored at -20°C and about 25% of each sample was later used for data acquisition.

### Proteomics Data Acquisition

For HeLa standard measurements, samples were diluted with Buffer A (0.1% FA in ddH_2_O) to enable the injection of 1 μg in 10 μL volume. Peptides were separated using the Easy NanoLC 1200 fitted with a trapping (Acclaim PepMap C18, 5 μm, 100 Å, 100 μm × 2 cm) and an analytical column (Acclaim PepMap RSLC C18, 2 μm, 100 Å, 75 μm × 50 cm). The outlet of the analytical column was coupled directly to a Q-Exactive HF Orbitrap mass spectrometer (Thermo Fisher Scientific). Solvent A was 0.1% (v/v) FA, in ddH_2_O and solvent B was 80% ACN, 0.1% (v/v) FA, in ddH_2_O. The samples were loaded with a constant flow of solvent A at a maximum pressure of 800 bar, onto the trapping column. Peptides were eluted via the analytical column at a constant flow of 0.3 μL/minute at 55°C. During elution, the percentage of solvent B was increased linearly from 4 to 5% in 1 minute, then from 5% to 27% in 30 minutes, and then from 27% to 44% in a further 5 minutes. Finally, the gradient was finished with 10.1 minutes at 95% solvent B, followed by 13.5 minutes at 96% solvent A. Peptides were introduced into the mass spectrometer via a Pico-Tip Emitter 360 μm OD × 20 μm ID; 10 μm tip (New Objective) and a spray voltage of 2 kV.

For FFPE lung adenocarcinoma measurements, about 25% of each sample was used for direct injection. Peptides were separated using the Easy NanoLC 1200 fitted with a trapping (Acclaim PepMap C18, 5 μm, 100 Å, 100 μm × 2 cm) and a self-packed analytical column (Reprosil-Pur Basic C18, 1.9 μm, 100 Å, 75 μm × 40 cm). The C18 material was packed into fused silica with an uncoated Pico-Tip Emitter with a 10 μm tip (New Objective) using a Nanobaum pressure bomb. The outlet of the analytical column was coupled directly to a Q-Exactive HF Orbitrap (Thermo Fisher Scientific) mass spectrometer. Solvent A was 0.1% (v/v) FA, in ddH_2_O and solvent B was 80% ACN, 0.1% (v/v) FA, in ddH_2_O. The samples were loaded with a constant flow of solvent A at a maximum pressure of 800 bar, onto the trapping column. Peptides were eluted via the analytical column at a constant flow of 0.3 μL/minute at 55°C. During the elution, the percentage of solvent B was increased in a linear fashion from 3 to 8% in 4 minutes, then from 8% to 10% in 2 minutes, then from 10% to 32% in a further 68 minutes, and then to 50% B in 12 minutes. Finally, the gradient was finished with 7 minutes at 100% solvent B, followed by 10 minutes 97% solvent A. Peptides were introduced into the mass spectrometer at a spray voltage of 2.5 kV.

In both settings, the capillary temperature was set at 275°C. Full scan MS spectra with mass range *m/z* 350 to 1500 were acquired in the Orbitrap with a resolution of 60,000 FWHM. The filling time was set to a maximum of 50 ms with an automatic gain control target of 3 × 10^6^ ions. The top 10 most abundant ions per full scan were selected for an MS^2^ acquisition. The dynamic exclusion list was with a maximum retention period of 60 seconds. Isotopes, unassigned charges, and charges of 1 and >8 were excluded. For MS^2^ scans, the resolution was set to 15,000 FWHM with automatic gain control of 5 × 10^4^ ions and maximum fill time of 50 ms. The isolation window was set to m/z 1.6, with a fixed first mass of m/z 120, and stepped collision energy of 28.

### Proteomics Data Processing

Raw files were processed using MaxQuant (version 1.5.1.2). The search was performed against the human Uniprot database (20170801_Uniprot_homo-sapiens_canonical_reviewed; 20214 entries) using the Andromeda search engine with the following search criteria: enzyme was set to trypsin/P with up to 2 missed cleavages. Carbamidomethylation (C) and oxidation (M) / acetylation (protein N-term) were selected as a fixed and variable modifications, respectively. First and second search peptide tolerance were set to 20 and 4.5 ppm, respectively. Protein quantification was performed using the label-free quantification (LFQ) algorithm of MaxQuant. LFQ intensities were calculated separately for different parameter groups using a minimum ratio count of 1, and minimum and average number of neighbors of 3 and 6, respectively. MS^2^ spectra were not required for the LFQ comparison. On top, intensity-based absolute quantification (iBAQ) intensities were calculated with a log fit enabled. Identification transfer between runs via the matching between runs algorithm was allowed with a match time window of 0.3 minutes. Peptide and protein hits were filtered at a false discovery rate of 1%, with a minimal peptide length of 7 amino acids. The reversed sequences of the target database were used as a decoy database. All remaining settings were set as default in MaxQuant. LFQ values were extracted from the protein groups table and log2-transformed for further analysis. No additional normalization steps were performed, as the resulting LFQ intensities are normalized by the MaxLFQ procedure. Proteins that were only identified by a modification site, the contaminants, as well as the reversed sequences were removed from the data set. All consecutive steps were performed in Microsoft Excel, Perseus (version 1.6.1.3), and the software environment R (version 3.5.1). The differential expression analysis of the ADC samples was performed using Limma moderated t-statistics (R package version 3.36.3)^27^. Here, the technical replicates and the patient-dependent batch effect were taken into account within the applied model. Proteins with a p-value lower than 0.05 and an absolute log2 fold change higher than 1 were considered as significantly changing. The resulting lists of significantly regulated proteins were subjected to a gene ontology (GO)-term enrichment analyses using the STRING: functional protein association network database^28^. The gene set enrichment analyses (GSEA) were performed using R package fgsea^29^ (version 1.6.0) with a p-value ranking of proteins, gene sets defined by the REACTOME pathway database (R package ReactomePA version 1.24.0)^30^, the minimum size of gene sets set to 15, the maximum size of gene sets set to 500, and the number of permutations set to 10000. The t-SNE analyses were performed using R package tsne^31^ (version 0.1-3) with a perplexity set to 2 and number of iterations set to 5000.

### Investigation of intra-day and inter-day Precision

To test the precision of SP3 sample handling, we followed guidelines of the European Pharmacopoeia and the European Medicines Agency for the number of replicates necessary to validate our method^32, 33^. Specifically, we validated automated SP3 by an intra-day and inter-day component by processing a total of six 96-well plates with 10 μg protein of a HeLa batch lysate in each well in the morning and in the afternoon of three different days, over a time span of roughly one month, resulting in a total of 575 individual samples. Five randomly picked samples per plate (10 samples per day) were selected for direct LCMS analysis on the day of sample generation and a second technical-repeat injection of all 30 samples in a single batch acquisition. The number of samples per plate to be analyzed was chosen as a fair compromise to determine the precision of our sample processing with a reasonable amount of data acquisition time. The selected samples allowed the evaluation of the inter-day precision and intra-day precision while taking different processing times, plates, and buffers into account (robustness). The second technical injection in one batch allowed to evaluate the influence of longitudinal MS performance. Lastly, for the comparison of manual SP3 sixteen times 10 μg protein of a HeLa batch lysate were processed manually at the bench.

### Lower-limit of Starting Material

To evaluate the lower limit of processing capabilities of the Bravo SP3 setup, we generated starting material dilution series as follows: A) a dilution series of our standard HeLa protein stock, ranging from 10 μg to ∼5 ng in 1:2 dilution steps (10 μg, 5 μg, 2.5 μg, 1.25 μg, ∼625 ng, ∼312 ng, ∼156 ng, ∼78 ng, ∼39 ng, ∼19 ng, ∼10 ng, and ∼5 ng). The dilution series was generated and processed in four replicates on the same 96-well plate (12 concentrations and n=4). B) a dilution series starting from small numbers of counted cells that were directly transferred to a 96-well plate, ranging from 10.000 down to 10 cells. The dilution series was generated and processed in two plates à four replicate series (7 concentrations and n=8). Here, the European Pharmacopoeia recommends a minimum of three concentrations à three replicates^32^. In addition, two empty control injections were performed upfront of the data acquisition of each dilution series. The dilution series were measured in blank-interspaced blocks from lowest to highest concentrated samples to avoid potential carry over between injections.

### Assessment of Cross-Contamination

To assess potential cross-contamination between samples, we processed 24 wells of 10 μg standard HeLa protein stock interspaced with 24 empty controls. Seven peptide-containing samples and eleven empty controls were randomly selected for direct LCMS analysis. The number of samples to be analyzed was chosen as a fair compromise to determine potential carry over between wells during our sample processing with a reasonable amount of data acquisition time.

## Results

### Establishing a Generic, Automated Proteomic Sample Preparation Pipeline

SP3 method is a fast and simple clean-up procedure for unbiased clean-up of proteins and peptides from a wide variety of sample types, and its foundation on para-magnetic bead technology should render it adaptable to robotic automation. Here, we implemented SP3 on the Agilent Bravo liquid handling system, which is widely available to many laboratories. Doing so required optimization of a number of steps, including the positioning of required consumables, reagent and waste volumes, as well as the Bravo accessories, such as magnet, shaker, and heating block to ensure accessibility of all the required components for each consecutive task, such as tips-on, liquid aspiration and dispensing, and tips-off, including the required volumes for reagents, buffers, or waste. An overview of the deck setup is shown in Figure 1. As a result, the available deck space and the range of motion of the Bravo pipetting head were exploited to fully automate the process starting from cell or tissue lysates to peptides ready for MS analysis for 96 samples simultaneously.

**Figure 1:**
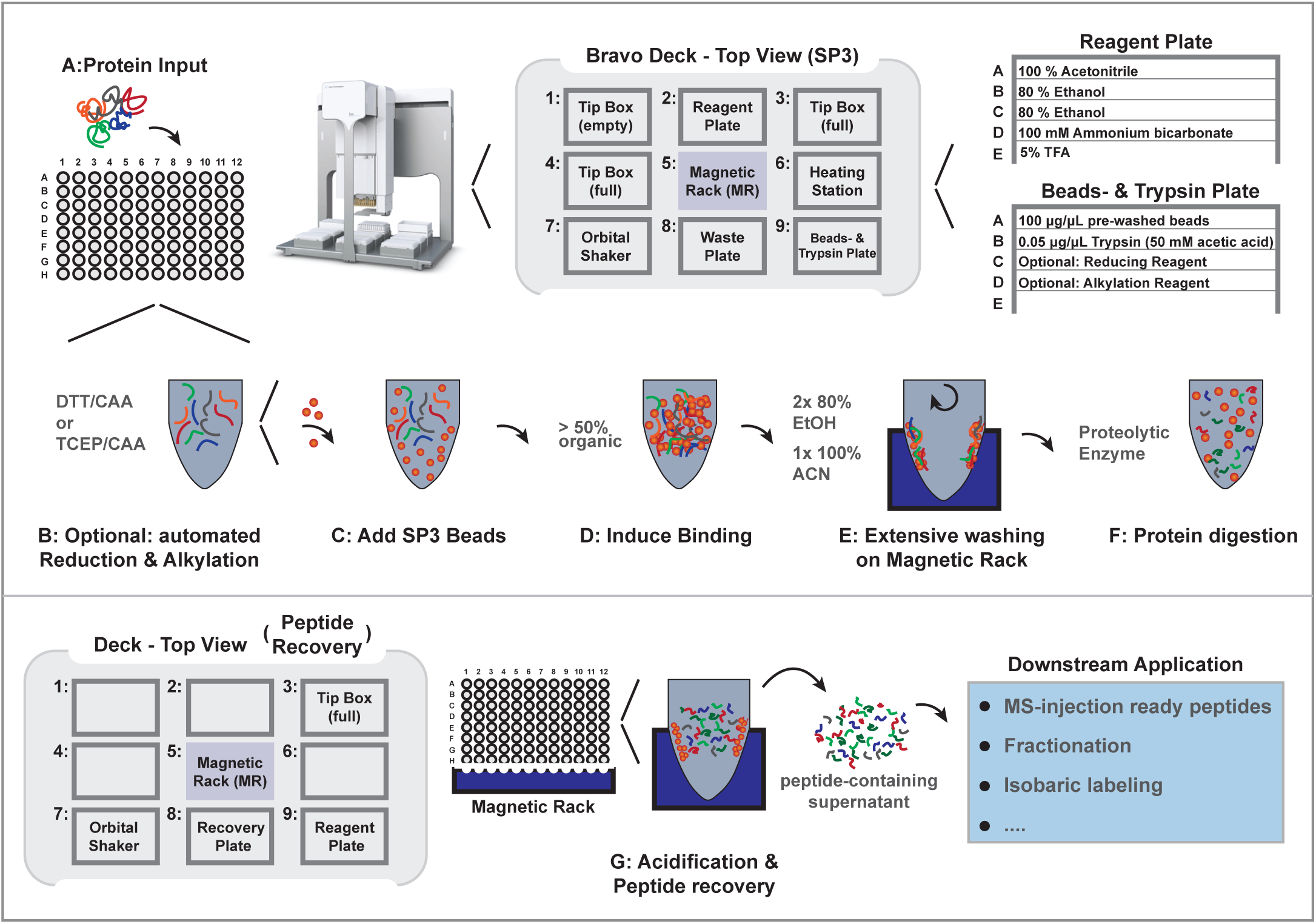
A schematic overview of automated single-pot solid-phase-enhanced sample preparation (autoSP3) workflow. The overview shows the different steps of the autoSP3 protocol from protein input to enzymatic digestion and peptide recovery. In addition, the setup of the Bravo deck is shown for the core clean-up protocol and separately for the peptide acidification and recovery step. The protocol ends with MS injection-ready peptide samples.

Initially, we implemented a procedure for protein reduction and alkylation using Dithiothreitol (DTT) & CAA (included in protocol B in **Supplementary Data B;** Supplementary Figure 1). However, to minimize the number of protocol tasks and simultaneously decrease the number of reagents, the protocol was adapted for a combined reduction and alkylation reaction with TCEP & CAA for 5 minutes at 95°C (included in protocol A in **Supplementary Data A**). Despite satisfying performance, the inefficient heating and cooling of the Bravo heating accessory, taking more than one hour to reach 95°C, led us to finally uncouple this step from the automated processing. Instead, proteins were manually reduced and alkylated using the combined reaction with TCEP & CAA for 5 minutes at 95°C in a PCR thermocycler (Protocol C & **Supplementary Data C**). Thereby, these three protocols (Supplementary Figure 1) leave various options open to the user, to either integrate reduction and alkylation with SP3 processing in a continuous but slightly longer procedure, or to perform this step off-deck in any preferred way to enhance speed and flexibility before transferring samples to the Bravo deck for automated SP3.

After manual or automated protein reduction and alkylation, the protocol continues (**Supplementary Protocol A or B**) or begins (**Supplementary Protocol C**) with the aliquoting of paramagnetic bead suspension to each sample (Figure 1A, 1B, and 1C). The liquid dispensing heights throughout the protocol were adjusted such that the pipette tips never contact the sample surface. From a suspension of paramagnetic beads (50 μg/μL in ddH_2_O), 5 μL are spotted as a droplet at the wall of each well and subsequently gently moved into the sample solution by agitation in an orbital shaking accessory. Next, ACN is added to each sample to a final concentration >50% to promote the trapping of proteins to the beads (Figure 1D). Here, ACN (Hughes et al., MSB 2014) is used rather than EtOH (Hughes et al., Nat Protocols 2019) because its pipetting properties are more reproducible without releasing hanging droplets at the end of each pipette tip. This is crucial, because organic solvent is added using a row of twelve pipette tips to dispense across the entire 96-well plate. Continuous switching between fast and slow agitation maintains a sufficient distribution of beads by preventing sedimentation and facilitating the efficient formation of protein-bead aggregates. Due to stickiness of beads in the presence of organic phase, we refrained from subsequent transfer or pipette mixing task to avoid loss of sample by beads adhering to the inner wall of the pipette tips. Since thorough mixing was shown as an essential part of effective washing and peptide recovery, the orbital shaking station was used to not only prevent sedimentation during the binding but also to improve the potency of each cleansing task. In brief, the rinsing of beads is performed as described in Hughes et al., Nat Protocols 2019^12^, with two times 80% EtOH and one time 100% ACN (Figure 1E). Between each of the washing steps, the supernatant is removed by trapping bead-bound proteins on a magnetic rack and removing the supernatant. The respective incubation times were evaluated and optimized to allow sufficient time for the beads to settle in a ring shape just above the bottom of the well before disposal of the wash solvent. The effective removal of wash solvents within each task is achieved by dividing each liquid aspiration task into two consecutive steps, in which the latter aspirates an additional air plug to avoid hanging droplets, as mentioned previously. Furthermore, specific liquid classes with optimized aspirating and dispensing velocities were defined for optimal movements (see protocol *.vzp files). Thus, the complete clearance of all residual solvents is achieved, for example, taking into account the propensity of ACN and especially EtOH to drain from the side wall in each well. This is especially important before adding trypsin to ensure a residual concentration of less than 5% ACN, corresponding to a maximum volume of 2 μL, which is compatible with protein digestion. Proteins trapped on the paramagnetic beads are resuspended in trypsin (or any other enzyme of choice) and incubated for two to 16 hours (Figure 1F). Following enzymatic digestion, peptide samples can be manually recovered and acidified in new plates or tubes (Figure 1G), or this can optionally be done on-deck after supplying new pipette tips (Protocol D & **Supplementary Data D**). During the automated SP3 protocol, both the paramagnetic bead stock as well as the enzyme solution (e.g. trypsin) and optionally reducing and alkylating reagents, are deposited in a second 96-well plate to ease pipetting of small volumes and to avoid uneconomical dead volumes of expensive reagents (Figure 1).

The autolysis of trypsin is prevented during the execution of the protocol by its storage in 50 mM acetic acid, and dilution with an adequate volume of 100 mM ABC at the time of mixing with the protein samples to achieve a digestion-compatible pH range.

In further optimization steps, the possibility of re-using tips for specific tasks was explored to increase sample throughput and reduce cost. Therefore, we adjusted the liquid dispensing heights in every task such that pipette tips never touch the sample surface or protein-bead aggregates. Thus, it was possible for every liquid-adding task to aspirate sufficient volume only once and successively dispense row-by-row across the entire 96-well plate. Subsequently, during any wash-disposal task, in which the pipette tips inevitably have to dip into the sample solution, aspiration velocities and heights were again optimized to allow liquid transfer without beads sticking to the pipette tips (see protocol *.vzp files). In addition, the same tip was re-used specifically for the same well in any liquid disposal task, thus excluding the risk of cross-contamination. To verify this experimentally, we processed 10 μg HeLa protein samples alternating with empty controls across half a 96-well plate. A subset of seven peptide-containing samples and eleven empty controls were randomly selected and subjected to direct LCMS data acquisition, as shown in Supplementary Figure 2A and 2B. Compared to MS intensities in sample-containing injections, most of the empty injections had a residual intensity of less than 0.03%, and in all cases well under 1% (Supplementary Figure 2B). These residual intensities could be primarily attributed to autolytic peptides of trypsin (which was added to all samples, including empty ones), and to (non-peptidic) contaminants with a +1 charge state, sharply contrasting with rich chromatograms from protein-containing samples (Supplementary Figure 2C).

**Figure 2:**
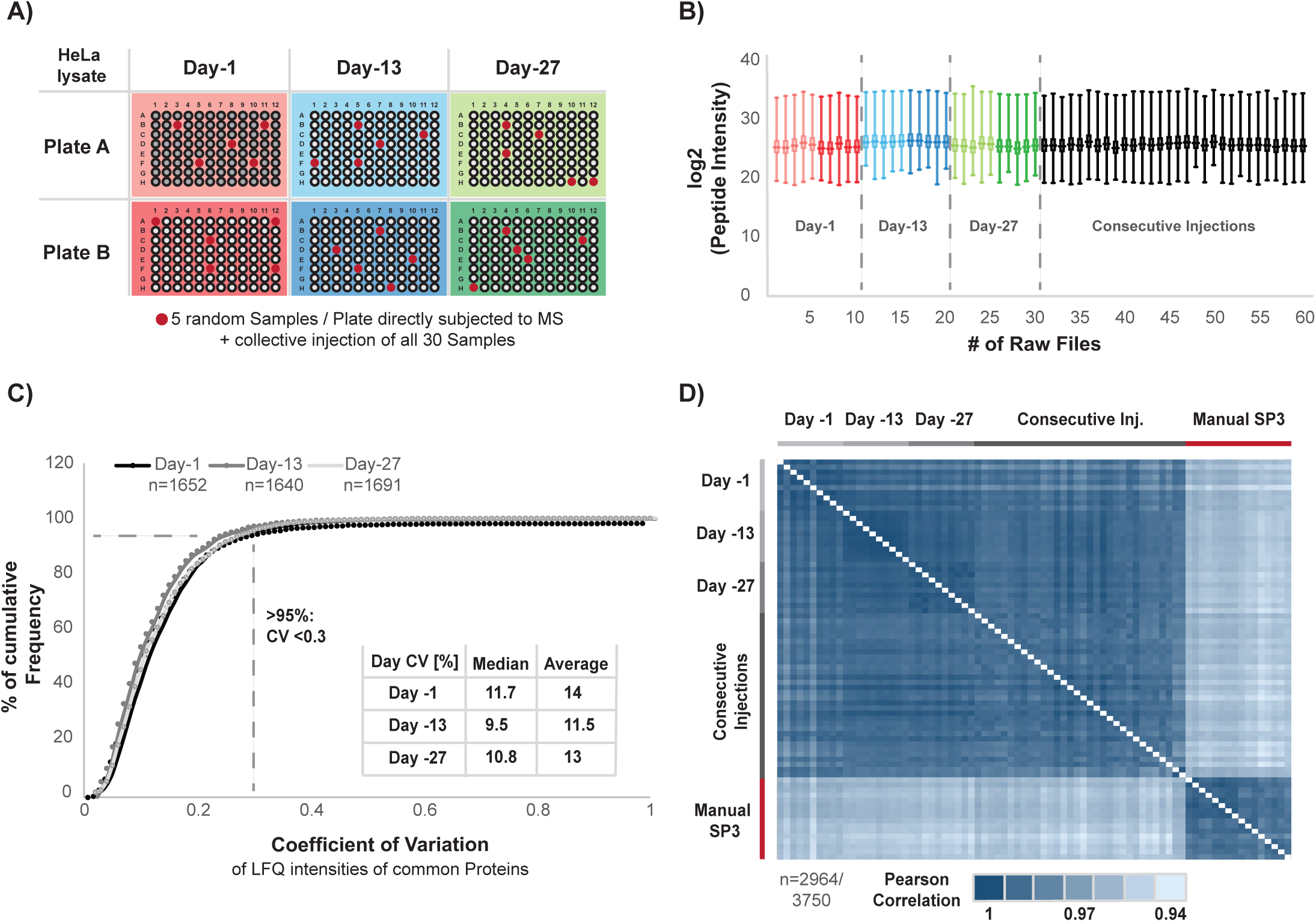
Evaluation of Intra-day and Inter-day Precision. A) a schematic representation of the experimental design. 96 times 10 μg protein of a HeLa batch lysate were processed in the morning (Plate A) and in the afternoon (Plate B) at three different days (Day-1, Day-13, and Day-27) over a period of a month. From each plate five randomly selected samples were subjected to direct LCMS analysis. In addition, all 30 samples (ten per day) were measured in a single combined batch to judge the influence of MS variability B) Box-whisker plots of log2 transformed peptide intensities across all 60 raw files. The color coding highlights the samples plate of origin. C) Cumulative frequency curve of the observed coefficient of variation (CV) of proteins that have been consistently identified and quantified within each day. Here, the ten raw files of each day are evaluated individually. The resulting median and average CV for each day are shown. D) Pearson correlation heatmap of all 60 raw files and additional sixteen manually prepared HeLa SP3 samples.

In summary, we established and optimized the SP3 protocol on a Bravo liquid handling system, taking care of all sample handling steps starting from 96 cell or tissue lysates and producing peptides ready for analysis by LCMS. A complete run of the Bravo SP3 protocol takes 1 hour and 23 minutes for 96 samples (protocol C) with an additional 45 or 65 min for reduction and alkylation with TCEP/ CAA (Protocol A) or DTT/ CAA (protocol B), respectively. The peptide-containing supernatant can be recovered without further clean-up for direct LCMS data acquisition or any compatible downstream protocol, such as high pH fractionation or TMT labeling. This can be done using a separate acidification and peptide recovery protocol, which takes about 7.5 minutes to complete (protocol D). The estimated cost for processing of 96 samples including reduction, alkylation, and peptide recovery is 92.39 euros (<1 euro per sample), including magnetic beads (600 μL of 50 μg/ μL), trypsin (576 μL of 0.05 μg/ μL), reagent and waste plates, three PCR plates, three pipette tip boxes, and all other buffers/solvents.

### Precision of automated SP3

A distinguishing feature of any automated procedure is strict standardization leading to precise and reproducible workflows. We evaluated this for automated SP3 by assessing its precision (defined by the EMA as the variability observed within the same laboratory^33^), both within the time span of 1 day (intra-day precision) as well as longitudinally over the period of 1 month (inter-day precision). To this end, HeLa cells were lysed, DNA and RNA was digested, and proteins were reduced and alkylated before transfer of protein to a 96-well plate for automated SP3 clean-up and digestion as described above. Intra-day precision was assessed by processing 96 times 10 μg protein of a HeLa lysate in the morning and in the afternoon of three different days (day 1, 13 and 27), i.e. over a time span of roughly one month, resulting in a total of six 96-well plates and 576 individual samples (Figure 2A). Inter-day precision was inferred by correlating data obtained across the three days. Specifically, LCMS was performed on the respective days immediately after completing automated SP3, by randomly selecting five samples from each of the six plates (i.e. 30 samples). In addition, we analyzed the exact same samples in one complete batch, resulting in additional 30 sample injections, to distinguish potential variance as a result of the SP3 processing from fluctuation in longitudinal MS performance. Collectively, this allowed us to evaluate the variability within a 96-well plate, within a day, across several days, as well as with and without potential variation in MS performance. Intensities of identified peptides were highly consistent across all samples with an average Pearson correlation of 0.9 between each sample, both within and across days (Figure 2B).

To assess intra-day precision at the protein level, we filtered the data obtained from the ten samples generated on a single day for proteins that had been identified and quantified without missing values, resulting in 1652, 1640, and 1691 proteins with an LFQ value for day-1, day-13, and day-27, respectively (Figure 2C). For each protein, the coefficient of variation (CV) across ten samples was calculated, demonstrating that more than 95% of the proteins quantified within each day had a CV of less than 30%, with a median CV of 11.7%, 9.5%, and 10.8% for day-1, day-13, and day-27, respectively, reflecting highly consistent protein quantification across replicates of sequentially processed sample plates and within days.

Next, we aimed to estimate inter-day precision of SP3 performance across all 60 data sets, and compare this to data from 16 samples prepared by manual SP3. To maximize the number of proteins included in this assessment, we applied the match-between-runs functionality in MaxQuant, increasing the proportion of proteins without missing values from 33.62% to 58.37%. An average of 14140 peptide spectrum matches and 3191 proteins (Supplementary Figure 3) were quantified with CVs of 7.1% and 2.1%, respectively. The median and average CV’s of the complete list of quantified proteins (n=3750) across all 60 samples was observed to be 18.1% and 20.5%. For further evaluation we selected proteins with an LFQ intensity in at least 45 out of 60 files, i.e. with a minimum data completeness of 75% across all measurements, corresponding to 2964 (79.04%) of the total number of quantified proteins, which were used for the subsequent comparison of variation within and across different days of sample preparation, as well as with and without the influence of daily MS performance (Table 1). CV’s across the 30 samples that were analyzed as one batch (median 14.3%, Table 1) were very similar to those analyzed immediately on the day of sample collection (median 13.3%, Table 1), indicating that differences in longitudinal MS performance were minimal, and that excellent CV’s can be obtained during sample and data acquisition over extended time periods. We observed excellent median and average CV’s of proteins across all 60 measurements (14.7% and 17.4%, respectively) showing a marginal but noticeable improvement as compared to the manually processed samples at (median CV 16.3%, average CV 18.6%; Table 1). This is further illustrated in Supplementary Figure 4A and 4B, showing consistently improved CV’s in automated vs. manual SP3 on a per-protein basis.

**Table 1.**
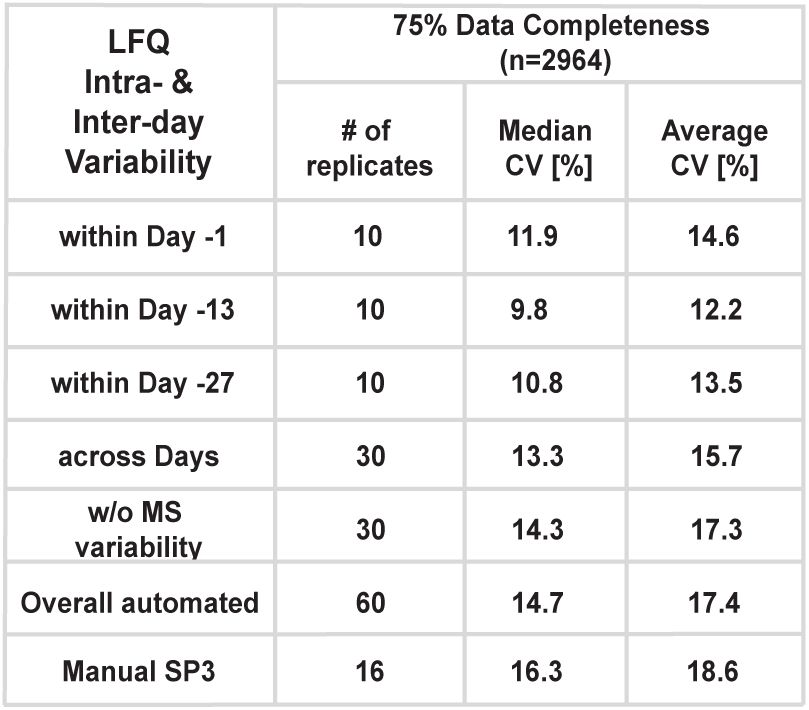
Summary of observed coefficient of variations (CVs). Corresponding to Figure 2, the table summarizes median and average coefficient of variation (CV) values for individual days, across days, with and without the MS imposed variability, and manual SP3.

Pearson coefficient between each of the 60 automatically and 16 manually prepared samples showed a very high correlation (>0.97) among both the 60 automatically and 16 manually processed samples, indicting highly robust performance by either procedure (Figure 2D). In addition, no differences are observable between data obtained on day 1, 13 and 27 indicating extremely high inter-day precision. Only slightly lower correlation (>0.94) was observed between data from manual or automated SP3, reflecting the high robustness of the SP3 protocol itself, but likely reflecting subtle differences between both protocols (e.g. sample volumes).

In addition, we sectioned all proteins identified across 60 experiments in four abundance bins (Figure 3A and Supplementary Figure 5), and investigate CV’s of their LFQ intensities. In the two highest abundance bins (A and B), >97.5% of the proteins have a CV <30%, with a median well under 10% (Figure 3A). In the lowest abundance bin (D) only 39.1% of proteins have a CV of less than 30%, which however comprises the group of ∼1000 proteins that were recovered by the match-between-runs option in MaxQuant. Without this, this number rises to 76.2%, (Supplementary Figure 5) while the median CV improves from 34.9% to 25%. Correspondingly, we looked at nine previously described housekeeping proteins^34^ and two randomly selected proteins with even lower abundance to check CV’s at the individual protein level. This demonstrated that CV’s both within and across days were well below 5% for the most abundant proteins and below 25% even for the low abundance proteins (Figure 3B, Table 2).

**Table 2.**
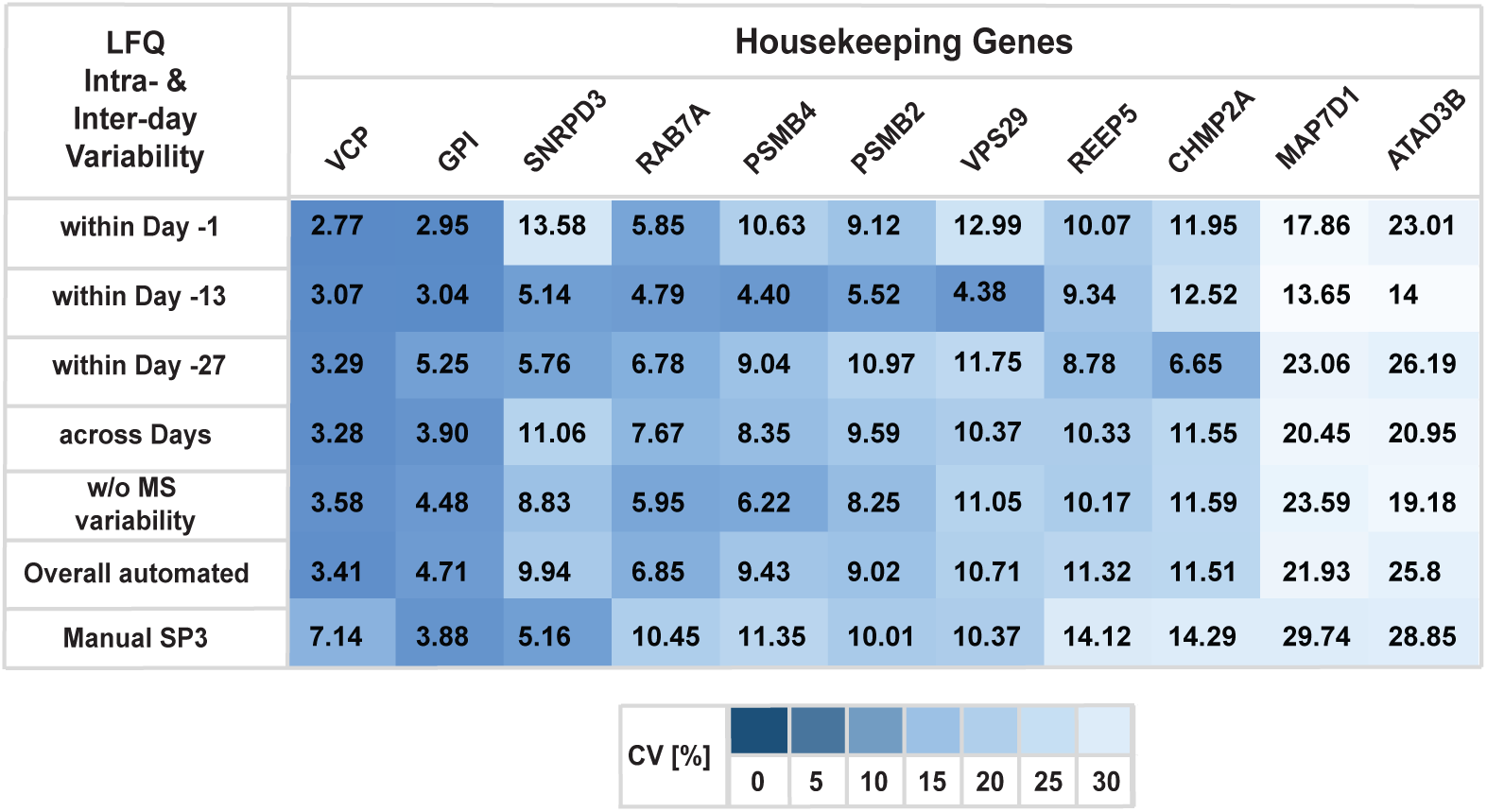
Summary of observed coefficient of variations (CVs). Corresponding to Figure 3B, the table summarizes average coefficient of variation (CV) values of individual selected proteins for individual days, across days, with and without the MS imposed variability, and manual SP3.

In conclusion, we have demonstrated robust performance of the SP3 method, irrespective of manual or automated processing, with slightly better median CVs for automated processing. This comes with additional benefits of high throughput, minimal hands-on time, and highly reproducible longitudinal performance over a period of several weeks.

**Figure 3:**
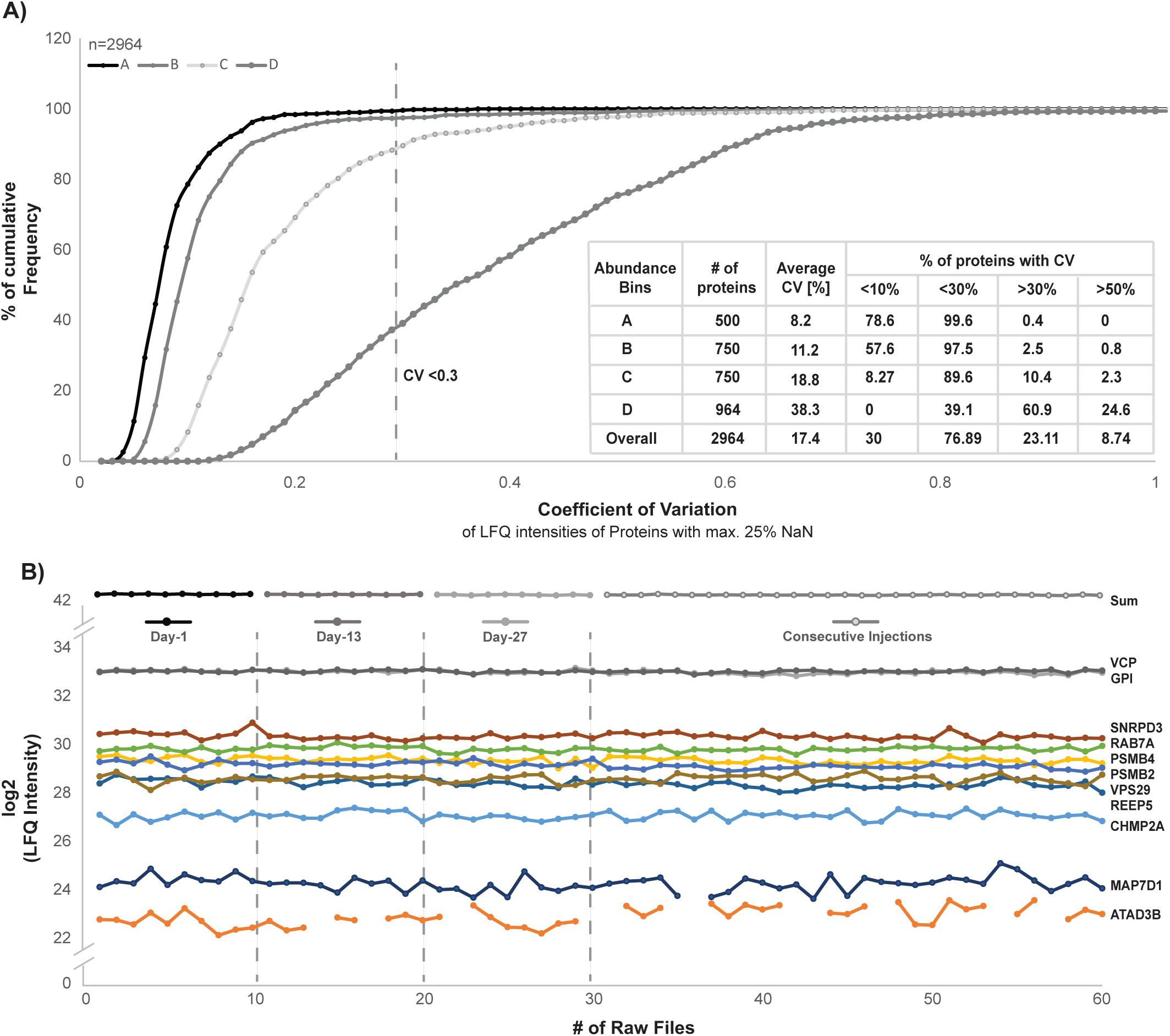
Protein abundance and coefficient of variation (CV). A) Four protein abundance bins (A, B, C, and D) were defined and cumulative frequency distributions of the calculated CVs of quantified proteins within each bin are plotted. The corresponding average CV values per group are shown. The table summarizes the percentage of quantified proteins observed with a CV higher or lower than 10%, 30%, and 50% for each abundance bin. B) log_2_ LFQ intensities of selected individual proteins and the sum of all proteins within a sample are plotted across all 60 measurements.

### Lower-Limit of Processing Capabilities of the Bravo SP3 Setup

A persistent challenge in proteomics is the consistent and sensitive analysis of low-input samples. Therefore, we aimed to investigate whether automated SP3 was capable of handling sub-microgram amounts of protein as input material, as we showed before for manual SP3 (Hughes et al 2014, Virant-Klun et al 2016)^11, 17^. Therefore, we prepared a 96-well plate with 4 replicates of 2-fold serial dilutions of a standard HeLa protein stock, ranging from 10 μg to ∼5 ng (**Figure 4A**), and processed them by automated SP3. The resulting 48 samples (twelve protein concentrations à four replicates) were injected for LCMS in blank-interspaced blocks of replicates from the lowest to the highest amount of protein. Entire samples were injected, except for the 4 highest concentrations which were maximized to 1 μg (back-calculated from the input) to avoid overloading of the analytical column. As expected, the number of quantified proteins and their summed intensities scaled with increased amounts of material, with narrow error distributions across the entire range indicating reproducible processing independent of input (**Figure 4A**). The 1 μg-injections from the four highest concentrated samples consistently identified ∼2000 proteins, indicative of high similarity in sample recovery off the magnetic beads independent of the amount of sample input. In addition, sub-microgram amounts of starting material were still sufficient to quantify several hundreds of proteins (e.g. 395 and 658 proteins from ∼39 ng and ∼80 ng, respectively). Strikingly, even ∼5 ng of protein input was sufficient to identify a median of 178 proteins (n=4) quantified with an LFQ value. This indicates very efficient protein capture, clean-up, and release by SP3, as well as digestion and transfer on-column with minimal protein loss.

**Figure 4:**
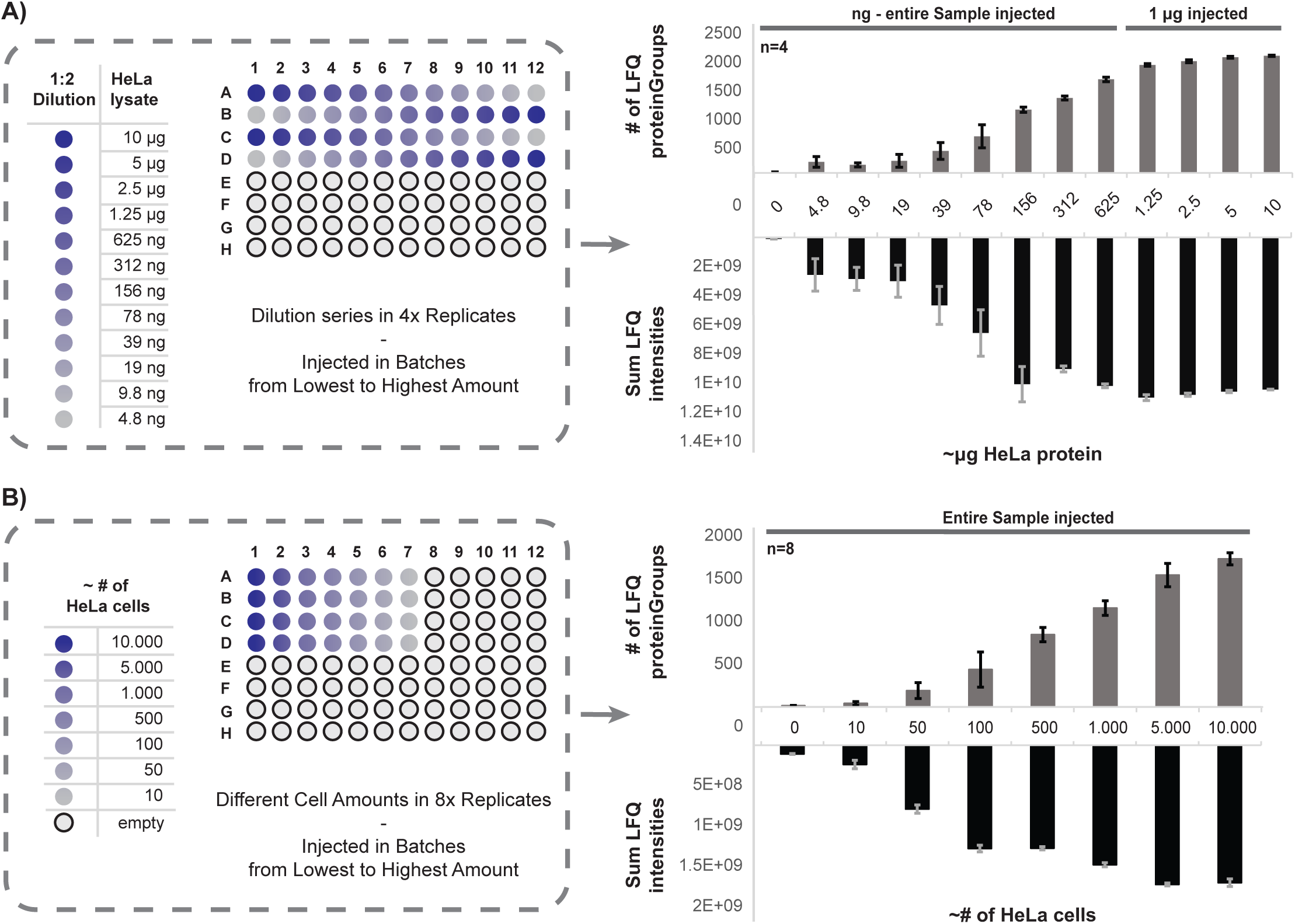
Evaluation of the lower-limit processing capabilities. A) Schematic representation of the experimental design with a 1:2 dilution series of a HeLa batch lysate starting from 10 μg down to 5 ng. The distribution of samples across the 96-well plate is shown. The dilution series was prepared in four replicates and samples were injected from lowest to highest concentrated. For the four highest concentrated samples 1 μg material was injected, whereas for sub-microgram samples the entire volume was used. The average number of quantified proteins per sample as well as the corresponding sum LFQ intensities are shown with error bars from the 4 replicates. B) Schematic representation of the experimental design of processing low numbers of HeLa cells. In total eight replicates of the following series of decreasing cell numbers were prepared “10.000, 5.000, 1.000, 500, 100, 50, and 10”. The average number of quantified proteins per sample as well as the corresponding sum LFQ intensities are shown with error bars from the eight replicates.

In another more realistic scenario of limited input material, we started from small numbers of cells instead of aliquoting from a common lysate. Therefore, HeLa cells were counted and directly transferred to a 96-well plate to create a range of samples containing 10,000 to 10 cells (in 8 replicates divided between 2 plates, **Figure 4B**), estimated to correspond to 1 μg to 1 ng of protein material (assuming 0.1 ng/cell). Cells were lysed in the plate, processed by automated SP3, and analyzed by LCMS. Again the number of proteins and LFQ values scaled with input, where the 1 μg-sample approached 2000 protein identifications (**Figure 4B**) as expected for this amount of input (compare to **Figure 4A**). The overall observed small error bars again demonstrate the reproducibility of the automated method and the capability of SP3 to process quantity-limited samples. Excitingly, the processing of as little as 100 cells was still sufficient to quantify on average 449 proteins (n=8).

In summary, the capability to reproducibly quantify 500-1000 proteins from an input of 100-1000 cells in a range below 100 ng input opens the door for multiple applications where sample availability is scarce, yet reaching sufficient depth for meaningful experiments. The ability to do so in an automated fashion removes the challenge of manual handling of such small samples.

### Application of autoSP3 to Clinical Pulmonary adenocarcinoma (ADC) Tumors

We next aimed to verify the performance of autoSP3 in a clinical real-world scenario processing a cohort of FFPE tissues, where SP3 is uniquely positioned to efficiently remove SDS used for de-crosslinking of this type of samples. Specifically, we aimed to understand the proteomic underpinnings of histological growth patterns observed in pulmonary adenocarcinoma. ADC is the most common histological lung cancer subtype accounting for roughly 60% of non-small cell lung cancers (NSCLC), known for their heterogeneous clinical, radiologic^35^, molecular^36–38^, and morphological^39^ features. Thus far, five distinct histological growth patterns have been recognized by the 2015 World Health Organization (WHO) Classification of Lung Tumors^25^. These growth patterns, which are being reported in any pathology report, have been proposed for tumor grading according to the predominant pattern of a tumor: lepidic (low grade; group 1), acinar and papillary (intermediate grade; group 2), and solid and micropapillary (high grade; group 3) (**Figure 5A**). Applying this grading system led to the observation of significant differences regarding prognosis^24^ and prediction of benefit from adjuvant chemotherapy^40^ where patients with lepidic ADC were associated with the most favorable and patients with micropapillary ADC with the worst prognosis. While marked gene expression differences have been identified for lepidic ADCs^41^, this is not the case for the other subtypes and thus requires further investigation to identify novel biomarkers, potential therapeutic targets, or to provide a functional explanation for the different growth patterns. In most invasive ADCs more than one growth pattern can be seen simultaneously, which further highlights the need to better understand functional differences and clinical implications of histological heterogeneity.

**Figure 5:**
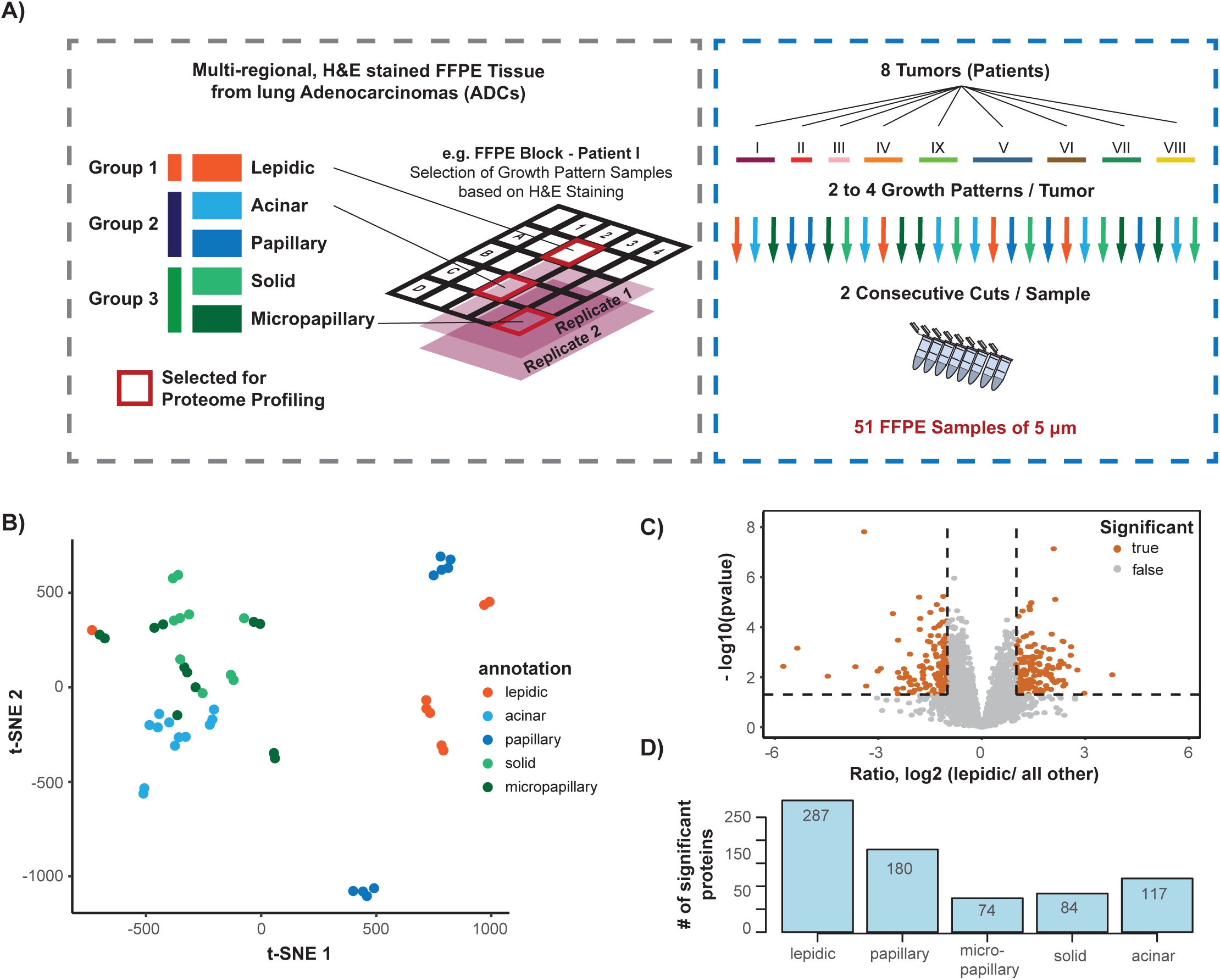
Application to clinical pulmonary Adenocarcinoma (ADC) FFPE tissue. A) Schematic illustration of the sample collection is shown. Samples were collected from eight different patient tumors. For each tumor, sections were processed with hematoxylin & eosin (H&E) staining to locate different growth patterns of lepidic (low-grade; group 1), acinar and papillary (intermediate grade; group 3), and solid and micropapillary (high-grade; group 3). Two to four growth patterns per tumor were selected and sectioned in two consecutive 5um iterations to provide replicates as close as possible, resulting in a total of 51 samples (one missing iteration). B) t-distributed stochastic neighbor embedding (t-SNE) analysis of the proteome data corrected using a linear regression model. The different growth patterns are highlighted and separated from each other. C) Volcano plot showing differential expression analysis using Limma moderated t-statistics for the comparison of lepidic samples against all others. Proteins passing significance thresholds of -log10 p-value < 0.05 and an absolute log_2_ fold change of 1 are highlighted in orange. D) Summary of significantly expressed proteins in the comparison of each growth pattern against all others.

Therefore, we collected FFPE tissue samples (5 mm × 5 mm × 5 μm) in a multiregional approach from central sections of eight ADC that had been histologically analyzed using hematoxylin and Eosin (H&E)-staining to locate and distinguish the different growth patterns. Two to four growth patterns were selected per tumor, and sections were performed in two consecutive iterations to provide replicates with highest possible similarity, resulting in a total of 51 samples (Figure 5A). The tissue was collected in PCR 8-stripes, lysed in two batches, and transferred to a 96-well plate in a randomized fashion for autoSP3 clean-up and protein digestion. Peptide samples were batch-randomized and analyzed by LCMS. Injecting ∼25% of each sample resulted in the identification of on average 3576 proteins (Supplementary Figure 6A) with normally distributed LFQ quantification as expected (Supplementary Figure 6B). A t-distributed stochastic neighbor embedding (t-SNE) analysis perfectly grouped replicate samples together (Supplementary Figure 6C), despite their random distribution on the 96-well plate during the processing by autoSP3, as well as batch-randomization during data acquisition, highlighting the reproducibility of the workflow.

Since samples grouped per patient (Supplementary Figure 6C) and not by growth pattern (Supplementary Figure 6D) we applied a linear regression model to reduce batch effects. As a result, a t-SNE analysis now separated the three superordinate groups (Figure 5B). Specifically, lepidic and papillary samples were now clearly separated from the remaining samples, while the replicate iterations were still clustered. In addition, acinar, solid, and micropapillary samples clustered more closely but tend to separate at the superordinate level (green and light blue; Figure 5B). The dissimilarity between lepidic and all other samples was expected and in line based on previous reports, while the division of papillary samples in two distinct subclusters separated from the rest of group 2 (acinar) was surprising (further discussed below).

To gain insight into the growth pattern-specific proteomes, we performed a Limma moderated t-statistics differential expression analysis between each growth pattern versus all combined other samples (Figure 5C, and Supplementary Figure 7). Lepidic tissue against all other samples showed the highest number of differentially expressed proteins (287 proteins, Figure 5D and Supplementary Figure 7) as expected from the t-SNE analysis (Figure 5B). A STRING-based gene ontology (GO)-term enrichment analysis of these proteins showed the enrichment of collagens among proteins that are more abundant in lepidic samples (Supplementary Figure 8A), reflecting different composition of the extracellular matrix. Collagens have previously been correlated with lung cancer growth, invasion and metastasis^42, 43^. Furthermore, we identified a cluster of mitochondrial ribosomal proteins (MRPs) which have been reported as a predictor for survival and progression, and thus as a potential prognostic biomarker in NSCLC^44^. In a GSEA of the same data we found *metabolism of polyamines* and *glucose metabolism* enriched in all others over lepidic (Supplementary Figure 9A). Greater capabilities of polyamine synthesis have previously been correlated with accelerated tumor spread and generally higher invasiveness^45^, while glucose absorption and metabolism towards anaerobic pathways are a reported key characteristic of the majority of NSCLC, strongly correlated with higher aggressiveness^46^. All of the above findings are in line with the known higher aggressiveness and worse prognosis of group 2/3 (intermediate & high grade) compared to group 1 (lepidic, low grade)^25^.

In the comparison of papillary versus all other samples, secretory and exocytosis-regulating proteins were identified among the significantly expressed (GO-term enrichment: Supplementary Figure 8B). Correspondingly, GSEA analysis (Supplementary Figure 9B), identified *Golgi-associated vesicle budding*, *intra-Golgi* and *Golgi-to-ER trafficking*, as well as *retrograde transport at the trans-Golgi network* among the top 10 significantly enriched terms pointing to an involvement of the secretory pathway. Altogether, this suggests extensive interaction with the environment in papillary-specific pathology. Interestingly, a relation of secreted proteins and NSCLC has been discussed previously^47^ however without differentiating between individual growth patterns. In addition, we found *mitogen-activated protein kinases (MAPK)*- and *non-canonical NF-kappaB-signaling* enriched. Both signal transduction pathways for cell survival and proliferation are common mechanisms to maintain oncogenic growth in NSCLC^48, 49^, however our data suggest that they are pronounced in papillary tumors.

We next followed up on the striking observation that papillary samples were separated in two subclusters based on proteome profiles (Figure 5B & Supplementary Figure 10A). To identify the molecular distinction between them, we subjected the 267 proteins that were differentially expressed between papillary_1 and papillary_2 to a GSEA (Supplementary Figure 10B). On the one hand this highlighted collagen-related and extracellular matrix gene sets enriched within Papillary_2 (Supplementary Figure 10C), possibly indicating differences in the microenvironment of each of the two papillary subclusters. On the other hand, processes related to mRNA nonsense-mediated decay and translation were enriched in the Papillary_1 cluster, indicating differences in the elimination of dysfunctional mRNAs in both sub-groups. Further analyses are needed to understand these phenomena in more detail.

For the remaining three comparisons, (i.e. acinar, solid, and micropapillary vs all others) many differentially expressed proteins were observed (Supplementary Figure 7), however no functional network could be retrieved, suggesting more subtle functional differences not represented by broad GO-categories. For example, we found increased expression of SCGB3A1 (log2 FC>2.5) and EEF1A2 (log2 FC>2.4) in micropapillary samples. SCGB3A1 has previously been found to be frequently methylated in NSCLC^50^ and was proposed as a novel tumor suppressor candidate in lung tumors. The methylation state was also significantly associated with the extent of the disease when comparing localized versus metastatic tumors. Furthermore, EEF1A2 has a canonical role in protein synthesis, cell proliferation and migration, and has been reported as a putative oncogene of NSCLC associated to poor prognosis showing higher expression in 28% of all cases^51^. In solid samples, we identified an upregulation (log2 FC>1.7) of MXRA5, which is aberrantly expressed in NSCLC and correlated to tumor progression and overall poor survival^52^. Its overexpression was used as an independent prognostic factor and proposed to potentially have value as a novel therapeutic target for the treatment of NSCLC^52^. Lastly, we found an upregulation (log2 FC>3.7) of OLFM4 in the comparison of acinar versus all others, which might play a role in lung tumorigenesis^53^. Altogether, we found several proteins that have previously been implicated in NSCLC, and that we can now associate with specific growth patterns.

In summary, these data demonstrate the applicability of autoSP3 pipeline to generate quantitative proteome profiles by processing a cohort of clinical ADC FFPE samples. The generated data illustrate high precision by tightly grouping of biological replicates from randomized samples, and the capability to process FFPE samples for the generation of relevant proteome data revealing differential expression between growth patterns.

## Discussion

In this work, we have presented the implementation of SP3 on a Bravo liquid handling platform for generic, reproducible and parallelized proteomic sample processing. Sample preparation is the only segment in the proteomic workflow that still largely relies on a series of manual handling and pipetting steps, including protein reduction/alkylation, clean-up to remove contaminating buffer components, and protein digestion. By seamlessly integrating all these steps into a fully automated process, autoSP3 alleviates many shortcomings that are associated with manual processing. Indeed, we demonstrate excellent reproducibility, shown in a series of 60 HeLa samples processed spread out over a month’s time (median CV of 16.3%), and in a cohort of 51 FFPE samples, where replicate tissue slices originating from subsequent cuts always grouped together despite randomized processing during autoSP3 and ensuing LCMS. The upshot of this is that samples can be generated over extended periods of time, e.g. during time series or longitudinal tissue collection, without introducing variability due to sample handling. All other benefits of manual SP3 also propagate in autoSP3, including handling of detergent-containing samples, high sensitivity, and low cost. The ability to handle detergent-containing samples, including SDS, is a great benefit over other methods such as iST, adding flexibility to the choice of protein extraction methods and enabling handling of sample types that depend on inclusion of SDS, such as for protein extraction from FFPE tissue, as shown by the analysis of ADC samples. The attribute of manual SP3 to perform well with low sample inputs was also demonstrated in autoSP3, showing reproducible identification of 500 proteins from sample amounts as small as 100 HeLa cells (Figure 4). This foreshadows powerful applications for the routine analysis of rare cell types, either in a basic-biological or clinical setting, where e.g. FACS-sorting followed by autoSP3 and LCMS can be integrated into a streamlined workflow with no other manual intervention than transferring a sample plate from one platform to the next. Importantly, since manual handling of minimal sample amounts is challenging, an automated workflow will reduce technical variability, instead allowing a more insightful focus on biological differences. Finally, autoSP3 is fast and affordable, taking 1.5 hours to complete 96 samples at a cost of <1 euro per sample. The same Bravo-platform can process one or more subsequent batches of 96 samples, thus easily reaching a capacity for several hundreds of samples per day. This should suffice even for very large-scale studies, considering that time for subsequent LCMS analysis will be the main limiting factor.

AutoSP3 is one of the first workflows offering a complete and universal solution for hands-free sample processing, implemented on the Bravo liquid handling system that is widely available in many genomics and biochemistry labs. To facilitate facile adoption, we provide all instrument *.vzp files for the core autoSP3 workflow (**Supplementary Data C**) and the extended versions that also include reduction and alkylation (**Supplementary Data A & B**), as well as for post-digestion acidification and peptide recovery (**Supplementary Data D**). While this should readily work for routine work-up of a wide diversity of sample types, all steps can be changed to the user’s needs (e.g. in case other proteases, reduction & alkylation reagents, digestion buffer, or different volumes are required). In addition, we expect that SP3 can also be readily implemented on other platforms, possibly benefiting from larger deck-size for increased capacity or extended functionality, e.g. to include peptide purification or TMT labeling if desired. In this respect, it is interesting to note that our initial protocol for peptide purification by SP3^11^ was recently implemented on an (Eppendorf) liquid handling system^54^. This can be an attractive solution for applications beyond protein expression profiling, e.g. to clean up post-translationally modified peptides after specific enrichment methods. Yet we like to state here that usually no peptide purification is needed after autoSP3, and that this was not used for any of the applications described here: contaminants are efficiently removed even when starting from 5% SDS as in the cases of FFPE samples allowing direct injection of digested peptides for LCMS. On the other side of the spectrum, MS-analysis of intact proteins purified by SP3 was recently shown^55^, opening the perspective that the use of autoSP3 might be extended to fit in a workflow for top-down proteomics.

The application of autoSP3 to a cohort of 51 ADC samples demonstrated the ability to process FFPE samples. Grouping of proteomes from subsequent tissue sections demonstrated high reproducibility of the workflow, while separation according to ADC growth pattern indicated profound molecular differences between pathological groups. In particular, in single-shot MS-analyses, we identified proteins associated with reduced invasion enriched in lepidic samples, as expected from pathology. This represented many more lepidic-specific proteins than previously suggested from gene expression profiles^41^, implying that i) not all gene expression differences propagated at the protein level, and ii) that instead proteome differences arise that do not result from mere gene expression changes. For example, Molina-Romero et al used microarray gene expression analysis to identify a list of 13 genes with specific differential expression in the lepidic histological growth pattern^41^. In our dataset, we quantified two proteins (CTPS1 and SNRNP40) corresponding to the list of 13 genes, but also identified an additional 285 differentially expressed proteins that could be of interest for follow-up studies for their potential use as biomarkers or therapeutic targets. Across all growth patterns, we identified a wide diversity of protein classes to be differently expressed, including secretory factors, metabolic proteins, signaling proteins, and transcription factors. This indicates that there is no singular cellular event that can readily explain microscopically observed differences, requiring more in-depth exploration to understand functional differences, and possibly the hierarchical order between them. Interestingly, we observed distinct sub-grouping of papillary samples, the full functional implications of which will need to be explored in more detail. In either case, our data may form the basis to unravel molecular (protein) markers that may aid more detailed subgrouping to complement pathological classification.

Altogether, autoSP3 is uniquely positioned as a key building block in a clinical pipeline for routine proteome profiling: i) It provides the ability to process any tissue type, including FFPE tissue. Since histology and WHO classification of tumors almost entirely relies on FFPE, autoSP3 now opens the potential for proteome profiling of samples that have been collected over decades; ii) rapid processing by autoSP3 contributes to fast turn-around times, e.g. to fit NSCLC international guidelines for genetic analysis (< 10 days)^56^; iii) low-input capabilities of autoSP3 allows analysis of small biopsies, possibly including those currently not accessible for proteomics, or reduction of biopsy size to achieve higher tumor cellularity and thus specificity of the assay; iv) robustness of the method minimizes technical variability; v) low cost.

In conclusion, autoSP3 removes manual sample handling from proteomic workflows, improving reproducibility and throughput of any proteomic sample type in a rapid, sensitive, and cost-effective manner. It seamlessly feeds into subsequent LCMS to constitute a highly standardized pipeline that should contribute to the identification of biological or clinical determinants in cohorts of dozens or hundreds of samples.

## Acknowledgments

This research project was supported by the Excellence Cluster CellNetworks to J.K. We thank Gertjan Kramer for helpful discussions, and the Tissue Bank of the National Center for Tumor Diseases for excellent technical assistance.

## Data availability

The mass spectrometry proteomics data have been deposited to the ProteomeXchange^57^ Consortium via the PRIDE^58^ partner repository with the dataset identifier PXD014556.

## Author contributions

T.M. and J.K. designed the study; T.M. performed research; T.M. and M.K. analyzed data; R.L. D.N., and A.S. provided reagents and samples; T.M. and J.K wrote the paper with input from all authors.

## Abbreviations

ABC: Ammonium bicarbonate
ACN: Acetonitrile
ADC: Adenocarcinoma
BCA: Bicinchoninic acid assay
CAA: Chloroacetamide
CV: Coefficient of variation
DMEM: Dulbecco’s modified Eagle’s medium
DTT: Dithiothreitol
EDTA: Ethylenediaminetetraacetic acid
EtOH: Ethanol
FA: Formic acid
FBS: Fetal Bovine serum
FFPE: Formalin-fixed and paraffin-embedded
FWHM: Full width half maximum
H&E: staining Hematoxylin and Eosin staining
iBAQ: Intensity-based absolute Quantification
LCMS: Liquid Chromatography coupled to Mass Spectrometry
LFQ: Label-free quantification
MS: Mass spectrometry
MS/MS: Tandem mass spectrometry
NSCLC: Non-small Cell Lung Cancer
PBS: Phosphate-buffered saline
PCR: Polymerase chain reaction
PIC: Protease-inhibitor cocktail
QE HF: Q-Exactive High Field Mass Spectrometer
RSLC: Rapid Separation Liquid Chromatography
SDS: Sodium dodecylsulfate
SP3: Single-pot solid-phase-enhanced sample preparation
TCEP: Tris(2-carboxyethyl)phosphine
TFA: Trifluoroacetic acid
TMT: Tandem mass-tag
t-SNE: t-distributed stochastic neighbor embedding
WHO: World Health Organization

---

**Supplementary Figure 1.**
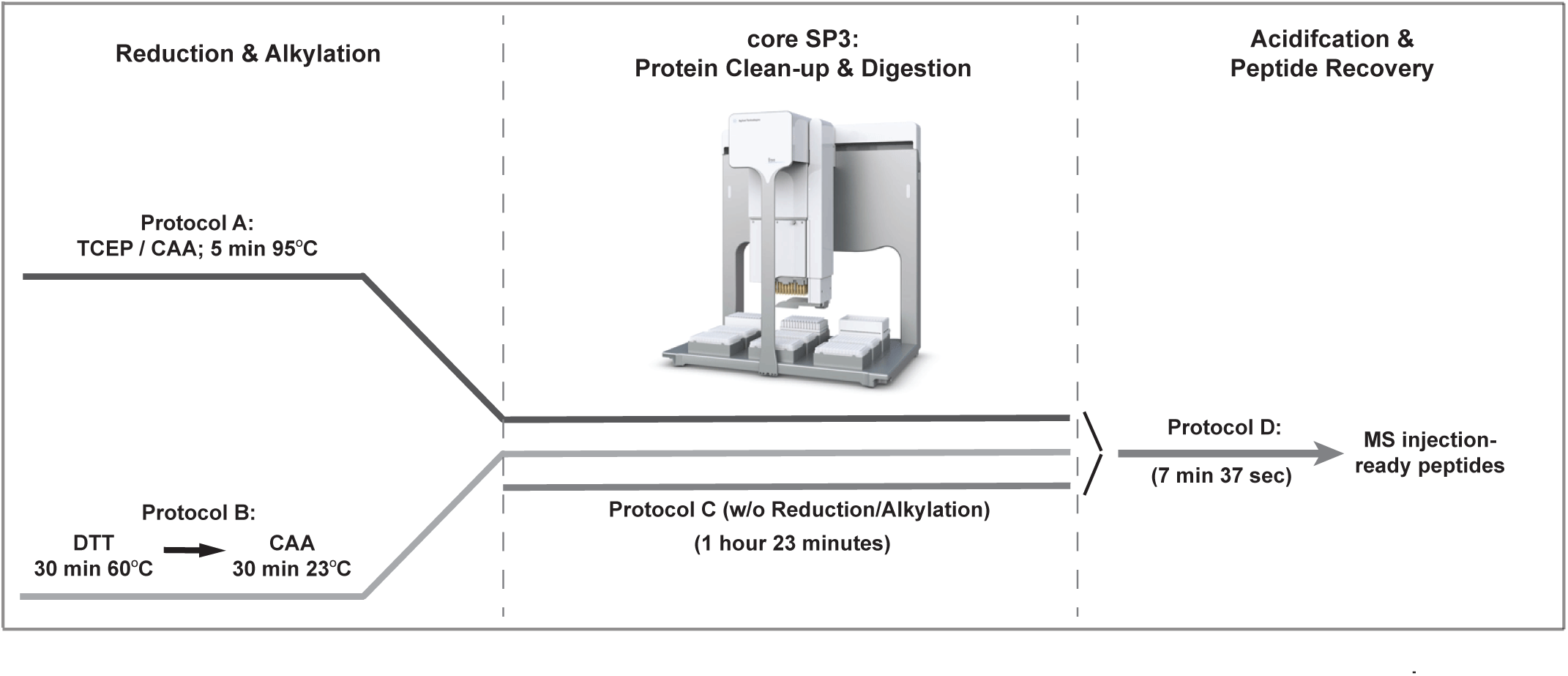
Schematic overview of available autoSP3 protocols. The autoSP3 protocol is provided with three different options for reduction and alkylation and with post-digestion peptide recovery as described in Supplementary Protocols (A to D). Protocol A: one-step reduction and alkylation using a TCEP/CAA mix for 5 minutes at 95°C, followed by SP3. Protocol B: two-step reduction and alkylation using DTT and CAA consecutively with 30 minutes incubation at 60°C and 23°C, respectively, followed by SP3. Protocol C: the core SP3 protocol omitting reduction and alkylation such that the user can flexibly process manually prepared samples. Protocol D: post-digestion acidification and MS injection-ready peptide recovery to a new sample plate.

**Supplementary Figure 2.**
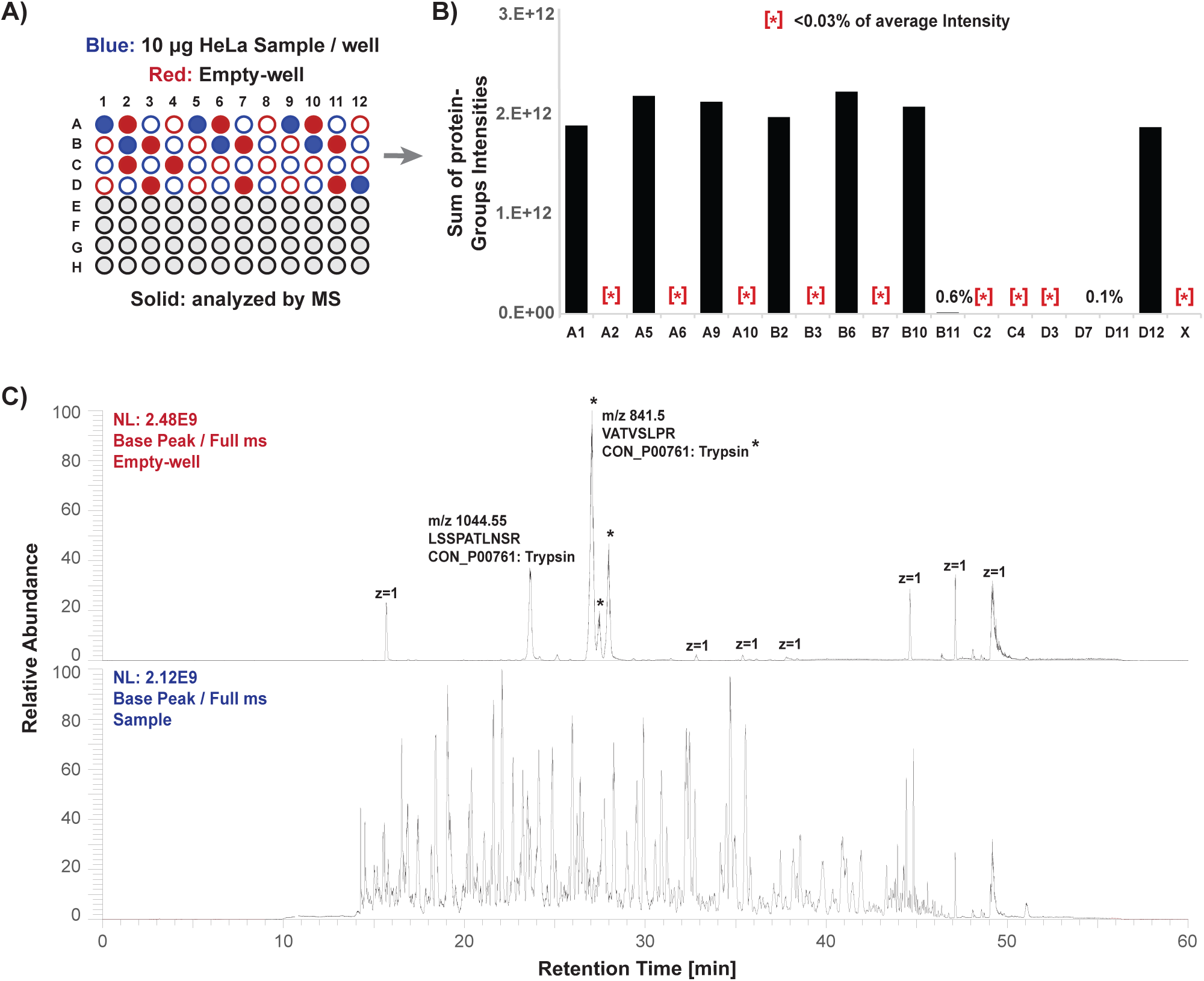
Experimental validation of absence of cross-contamination. A) Schematic representation of the experimental design to demonstrate the absence of cross contamination between wells. Half a plate (48 wells) was processed with 10 μg protein of a HeLa batch lysate in every second well (highlighted in blue) interspaced with empty wells as a control (highlighted in red). Randomly selected wells (highlighted in solid) were selected for direct LCMS. B) Bar plots of the summed intensities of protein groups across selected samples. A total of seven sample-containing injections were performed and a total of twelve empty controls. Asterisks indicate intensities <0.03%. C) Exemplary base peak MS1 spectrum for an empty control injection (top) and a sample-containing injection (bottom).

**Supplementary Figure 3.**
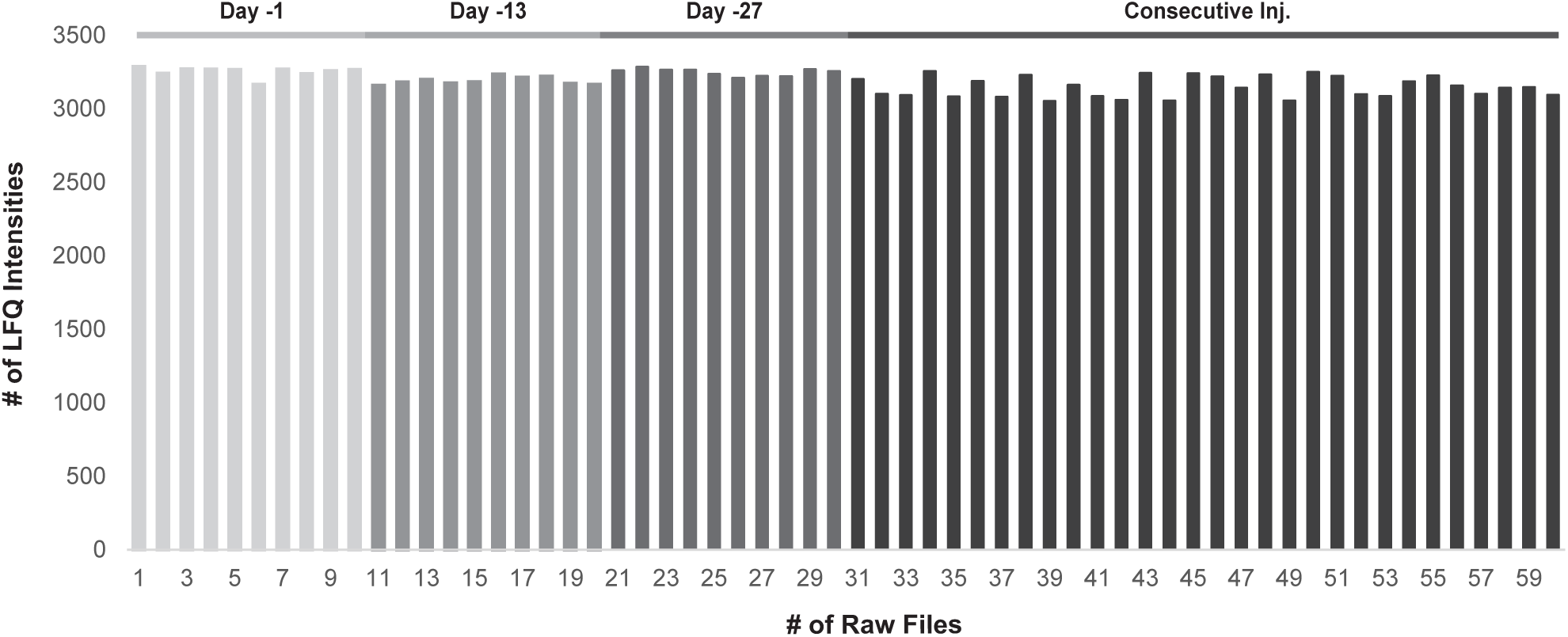
Protein identifications per sample. Bar plot summarizing the number of quantified LFQ protein groups per sample. Samples originating from different days and the consecutive injections of the same samples are highlighted in grey scales.

**Supplementary Figure 4.**
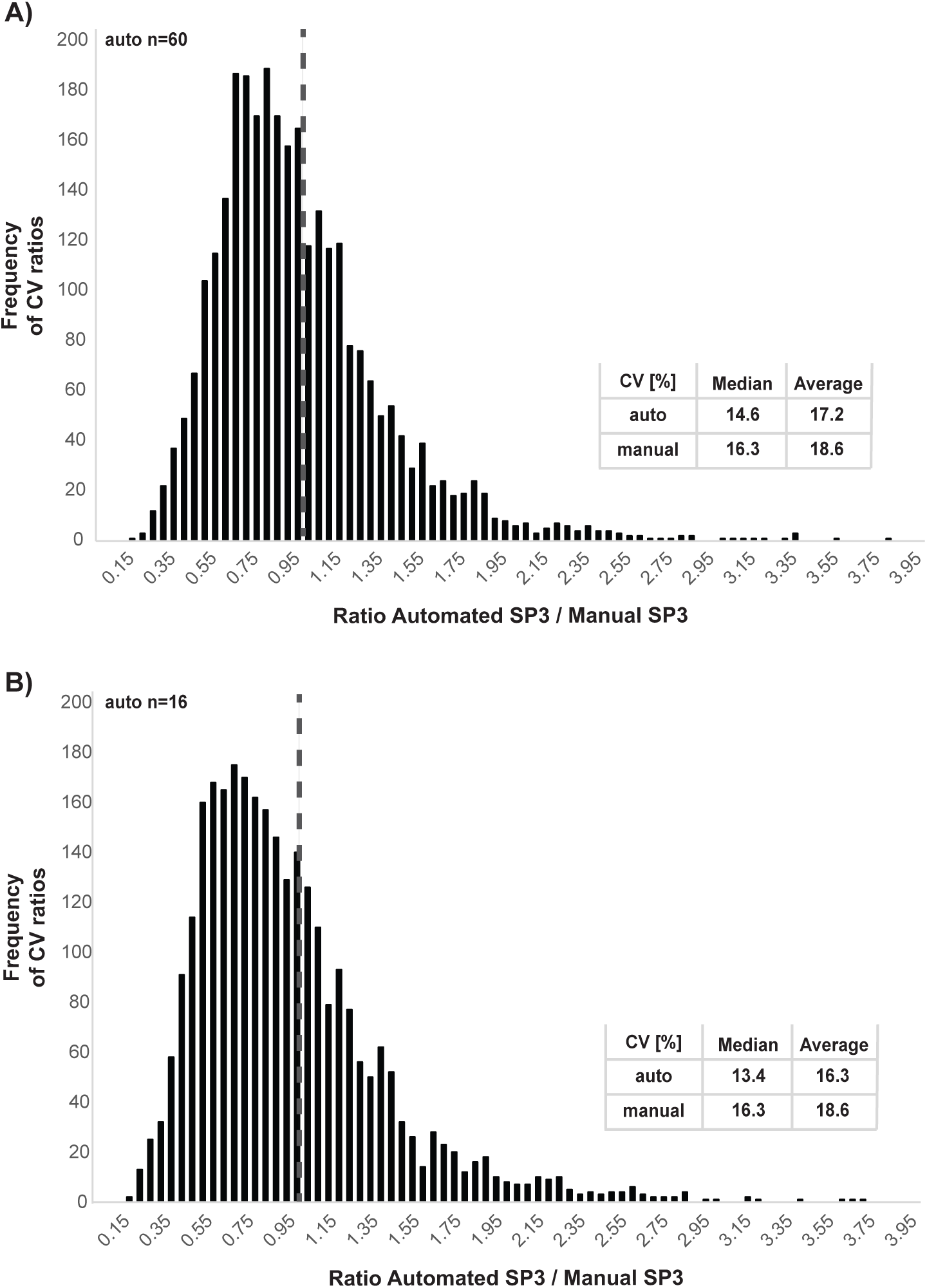
Comparison of coefficient of variation (CV) distribution between manually and automatically prepared samples. A) CVs of quantified proteins were calculated across all 60 acquired raw files and compared to sixteen manually prepared samples. A histogram of CV ratios of automatically prepared samples versus manually prepared samples is shown. The median and average CV is shown for both automatically and manually prepared samples. A dotted line highlights the ratio of 1. B) CVs of quantified proteins were calculated across sixteen randomly selected samples out of the 60 acquired raw files and compared to sixteen manually prepared samples. A histogram of CV ratios of automatically prepared samples versus manually prepared samples is shown. The median and average CV is shown for both, automatically and manually prepared samples. A dotted line highlights the ratio of 1.

**Supplementary Figure 5.**
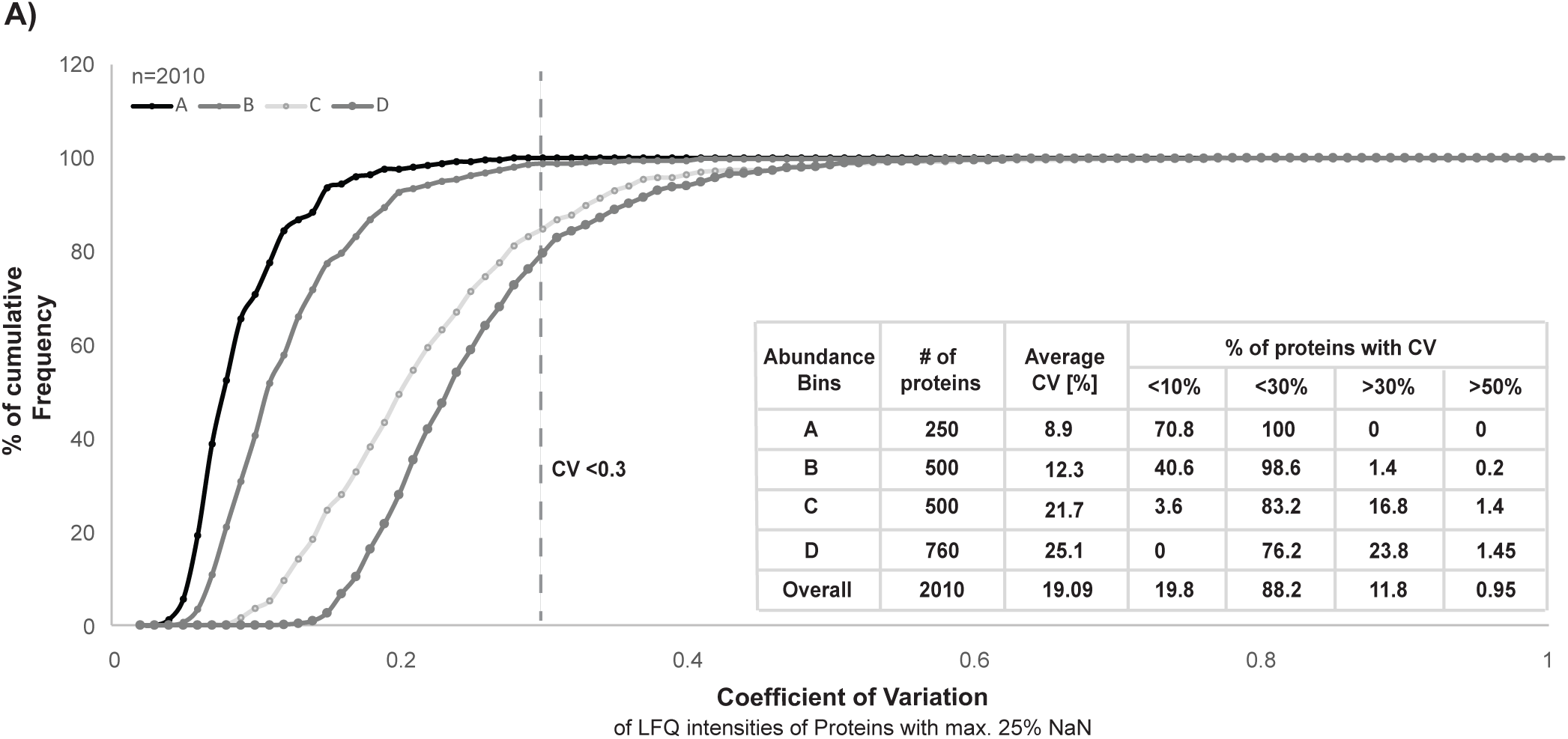
Protein abundance and coefficient of variation (CV) excluding matching between runs. Corresponding to Figure 3, four protein abundance bins (A, B, C, and D) were defined and cumulative frequency distributions of the calculated CVs of quantified proteins within each bin are plotted. The corresponding average CV values per group are shown. The table summarizes the percentage of quantified proteins observed with a CV higher or lower than 10%, 30%, and 50% for each abundance bin.

**Supplementary Figure 6.**
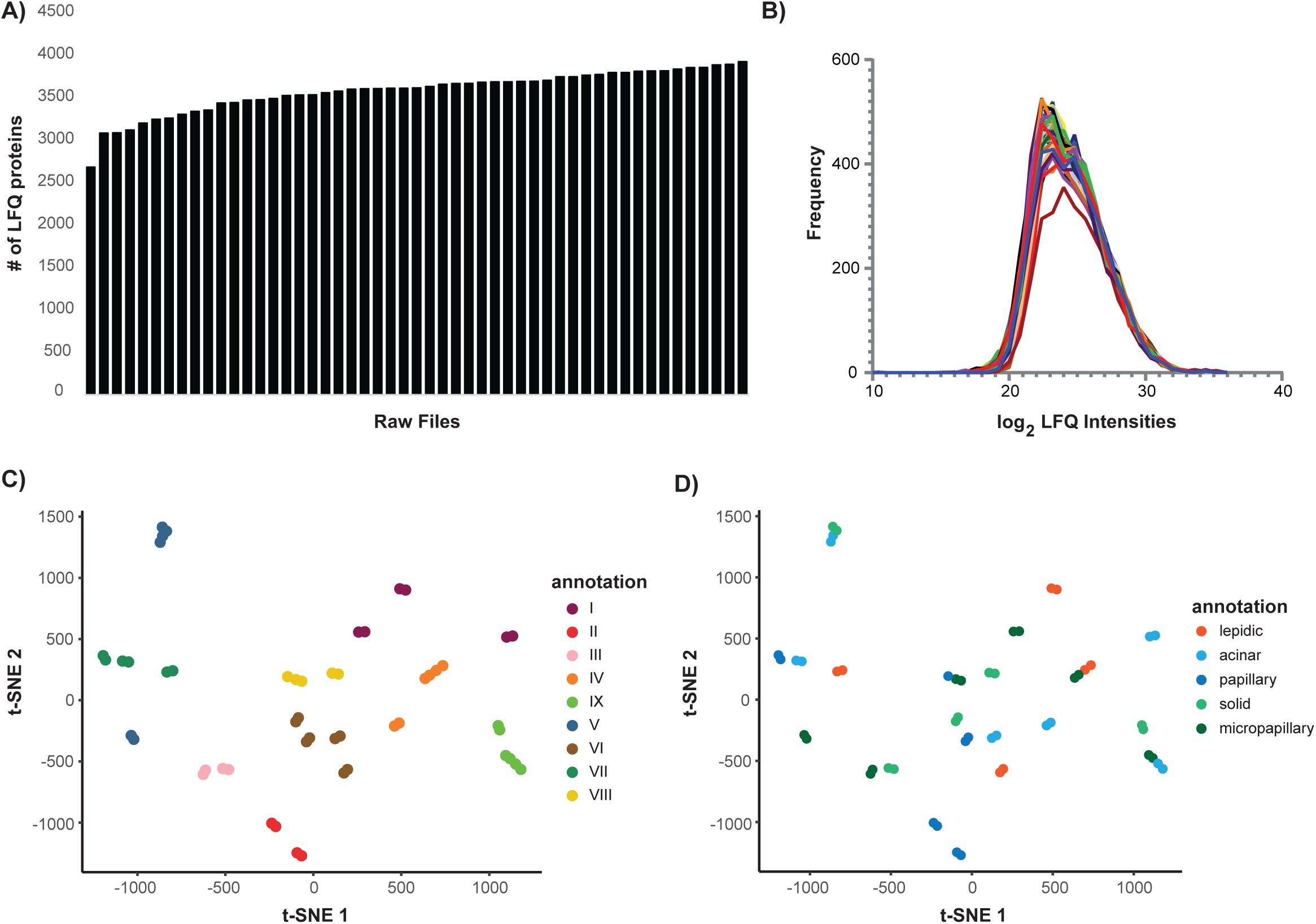
Identified and quantified proteins and uncorrected data of pulmonary adenocarcinoma (ADC) FFPE tissue. A) Bar plot summarizing the number of quantified LFQ protein groups per sample. B) Log_2_ LFQ intensity distribution for each sample highlights their normal distribution. C) t-distributed stochastic neighbor embedding (t-SNE) analysis of the uncorrected proteome data. The samples are highlighted according to their patient tumor of origin. D) Same as in C, highlighting the different growth patterns of each sample.

**Supplementary Figure 7.**
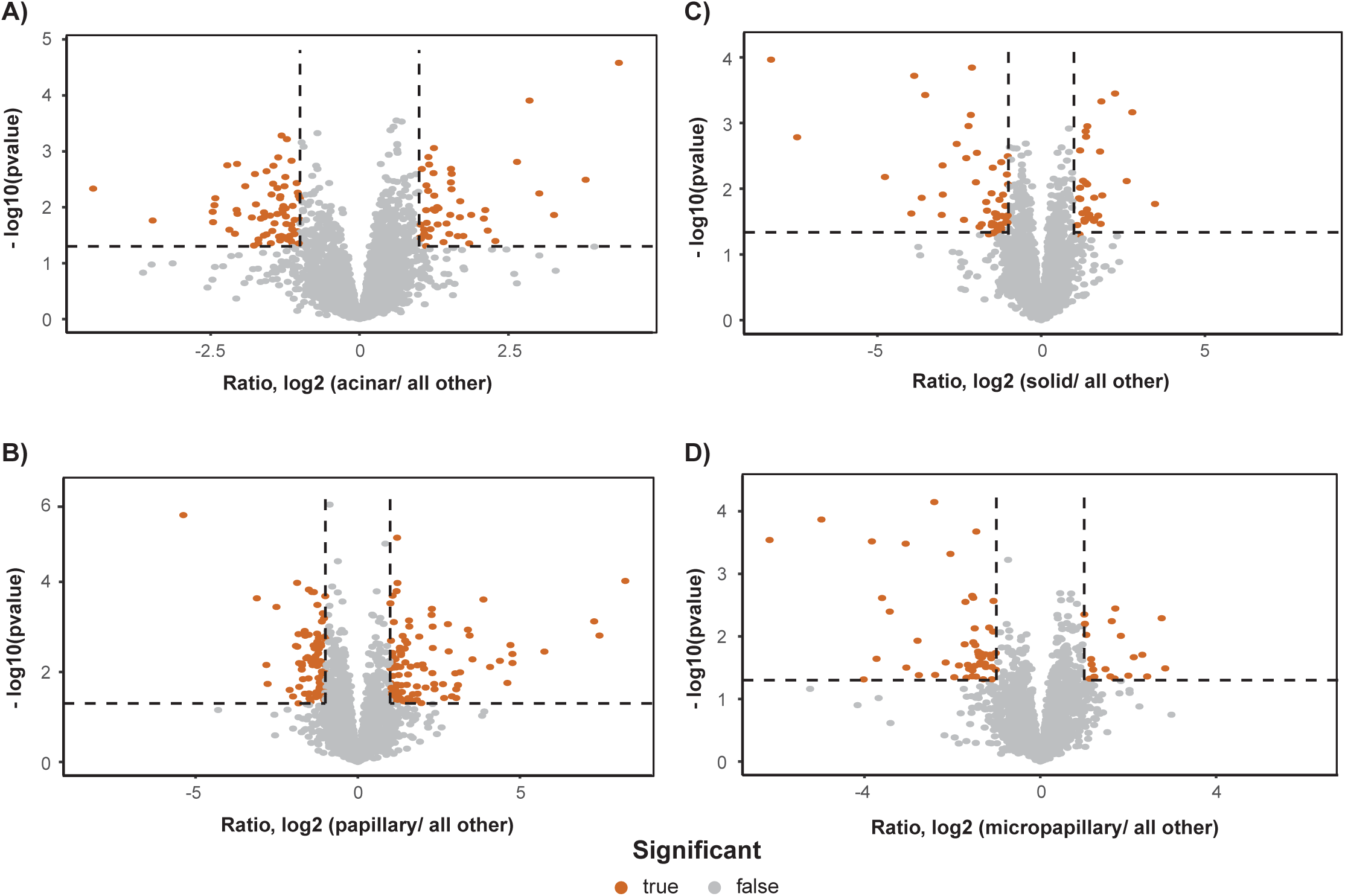
Differential expression analysis of remaining growth pattern comparisons. Corresponding to Figure 5, showing the differential expression analysis of acinar (A), papillary (B), solid (C), and micropapillary (D) tumors compared to all other samples by using Limma moderated t-statistics. Proteins passing significance thresholds of -log_10_ p-value < 0.05 and an absolute log_2_ fold change of 1 are highlighted in orange.

**Supplementary Figure 8.**
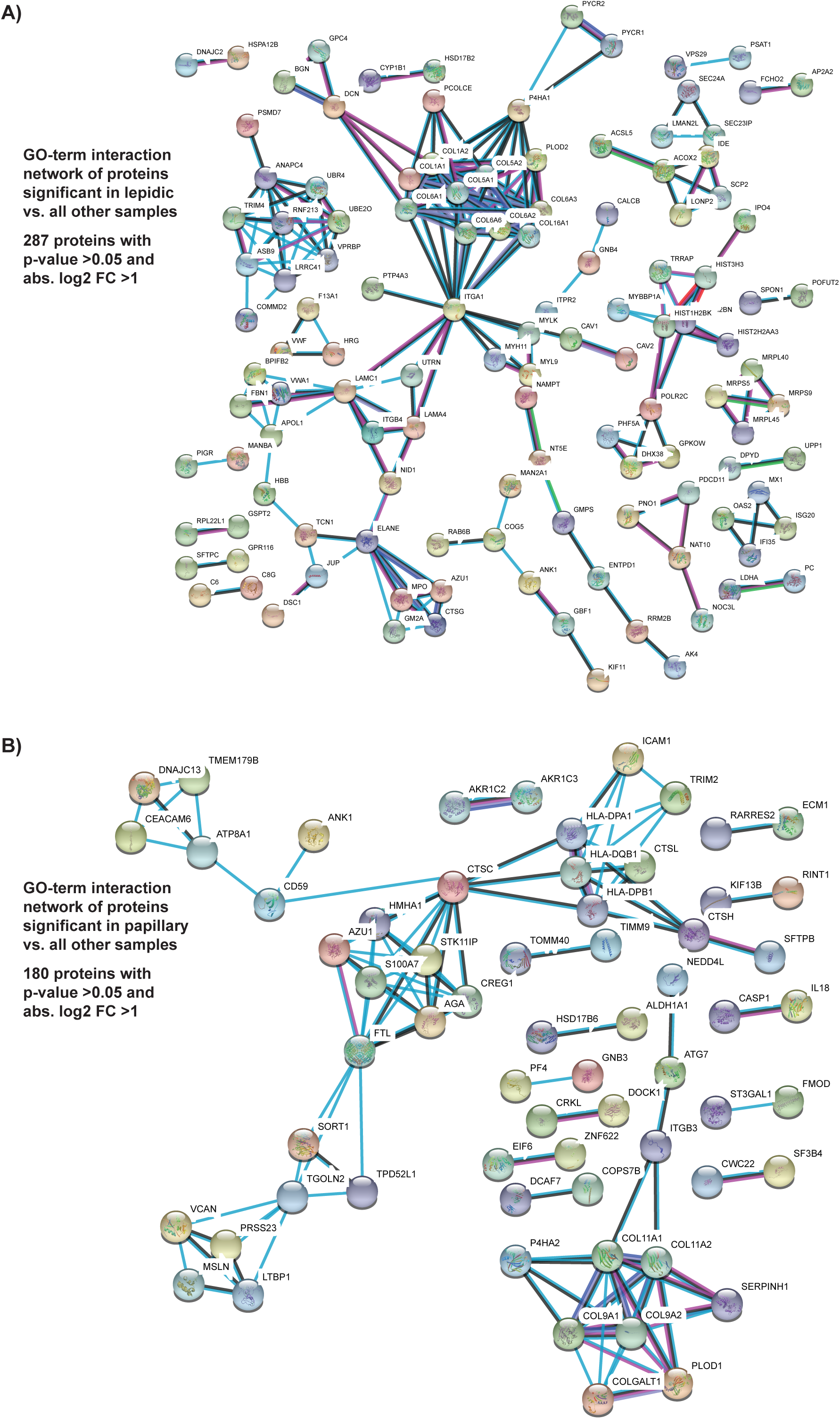
Gene ontology STRING interaction network analysis. A) STRING network analysis of the 287 significant proteins (-log_10_ p-value < 0.05 and an absolute log_2_ fold change of 1) in lepidic versus all other samples. B) STRING network analysis of the 180 significant proteins (-log_10_ p-value < 0.05 and an absolute log_2_ fold change of 1) in papillary versus all other samples.

**Supplementary Figure 9.**
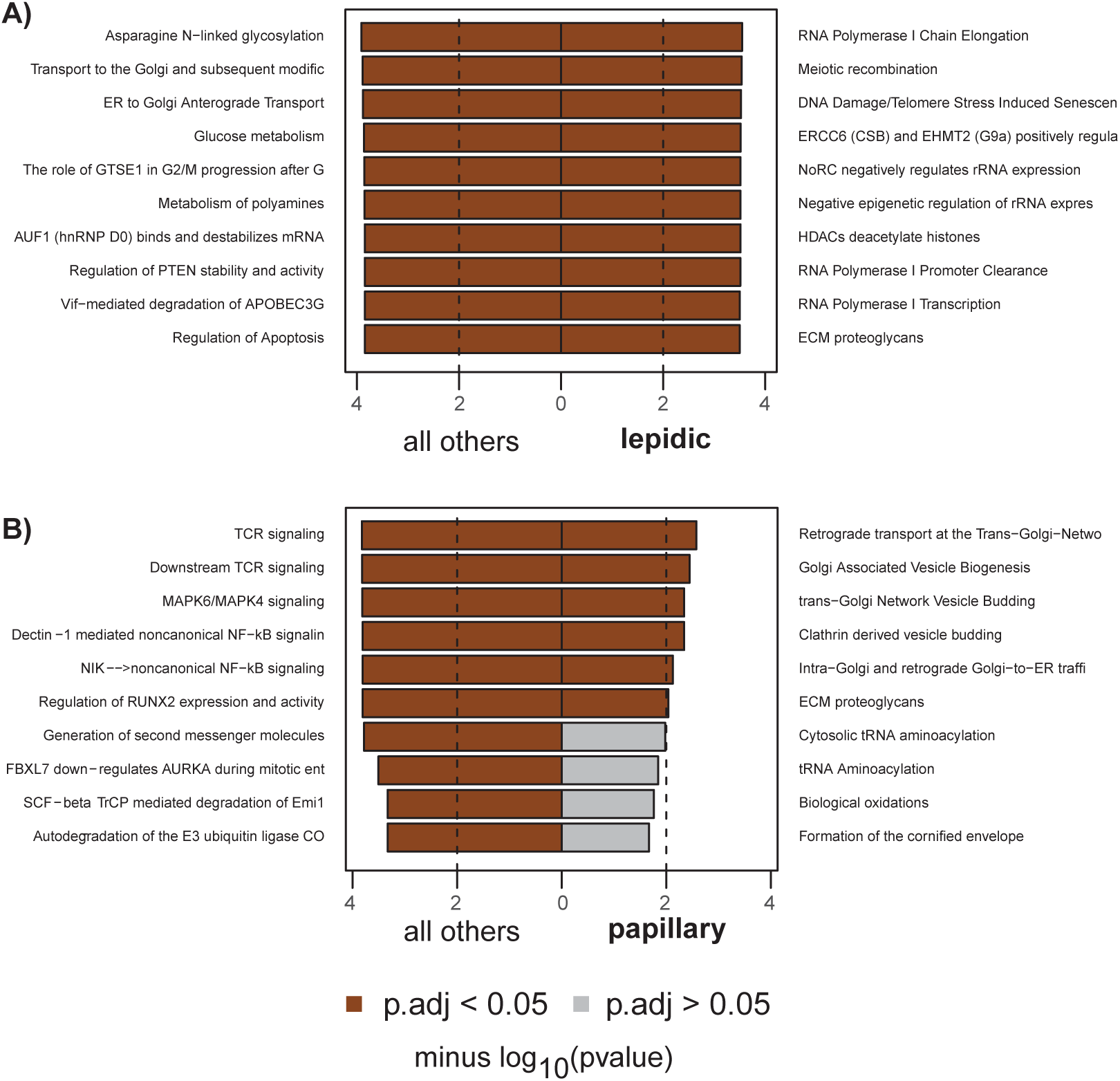
Gene set enrichment analysis (GSEA) of lepidic and papillary samples. A) Gene set enrichment analysis of p-value ranked proteins for lepidic versus all other samples. B) Gene set enrichment analysis of p-value ranked proteins for papillary versus all other samples. In both, gene sets with an adjusted -log_10_ p-value < 0.05 were considered significant and are highlighted in dark color.

**Supplementary Figure 10.**
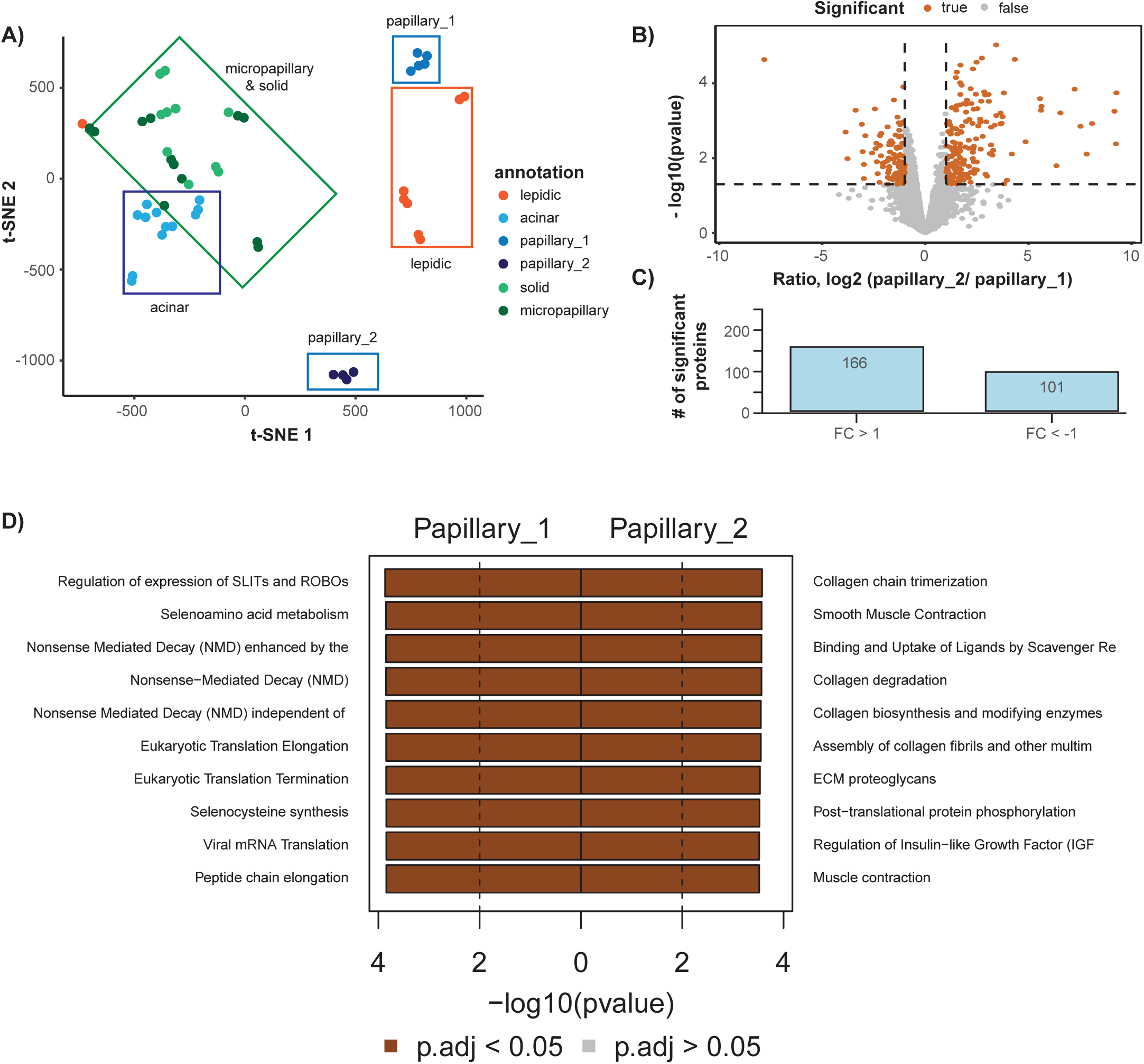
Differential expression analysis of papillary sub-groups. A) t-distributed stochastic neighbor embedding (t-SNE) analysis of the proteome data corrected using a linear regression model. The different growth patterns are highlighted according to their tumor cell content. B) Differential expression analysis between subclusters papillary_1 and papillary_2 (see A) using Limma moderated t-statistics. Proteins passing significance thresholds of -log_10_ p-value < 0.05 and an absolute log_2_ fold change of 1 are highlighted in orange. C) The number of differentially expressed proteins in the subcluster comparison. D) Gene set enrichment analysis of p-value ranked proteins for papillary_1 versus papillary_2. Gene sets with an adjusted -log_10_ p-value < 0.05 were considered significant and are highlighted in dark color.

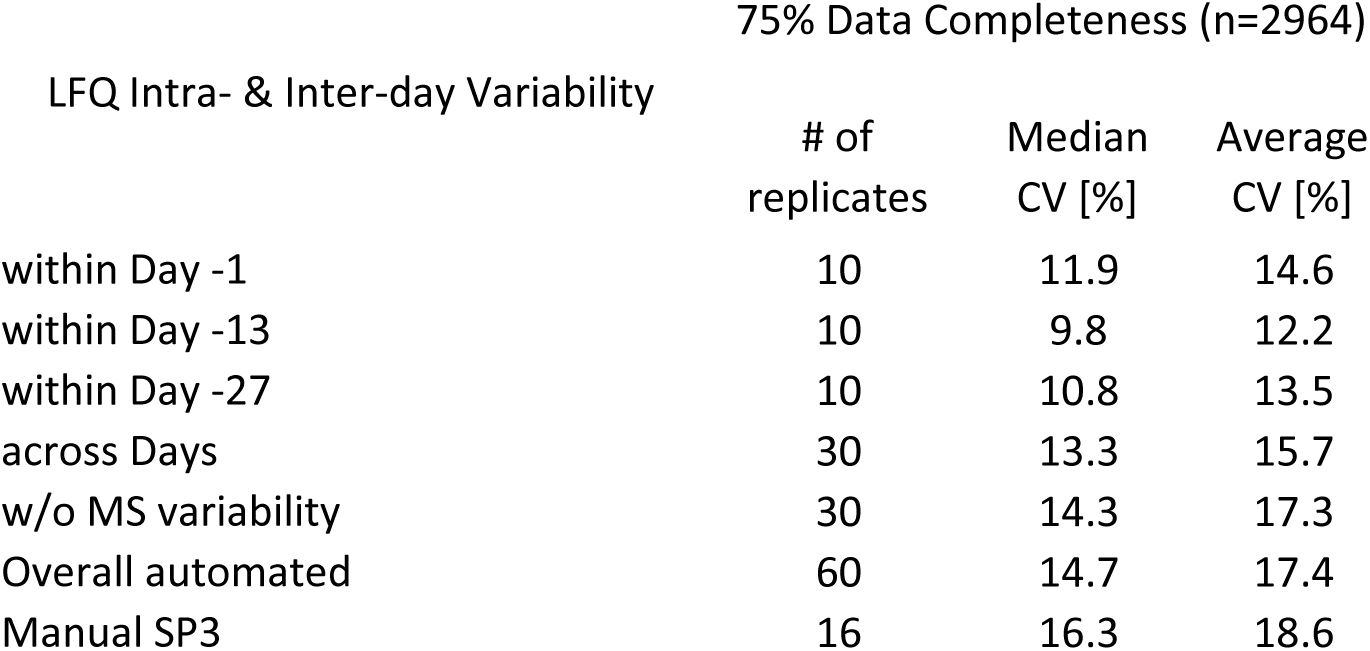

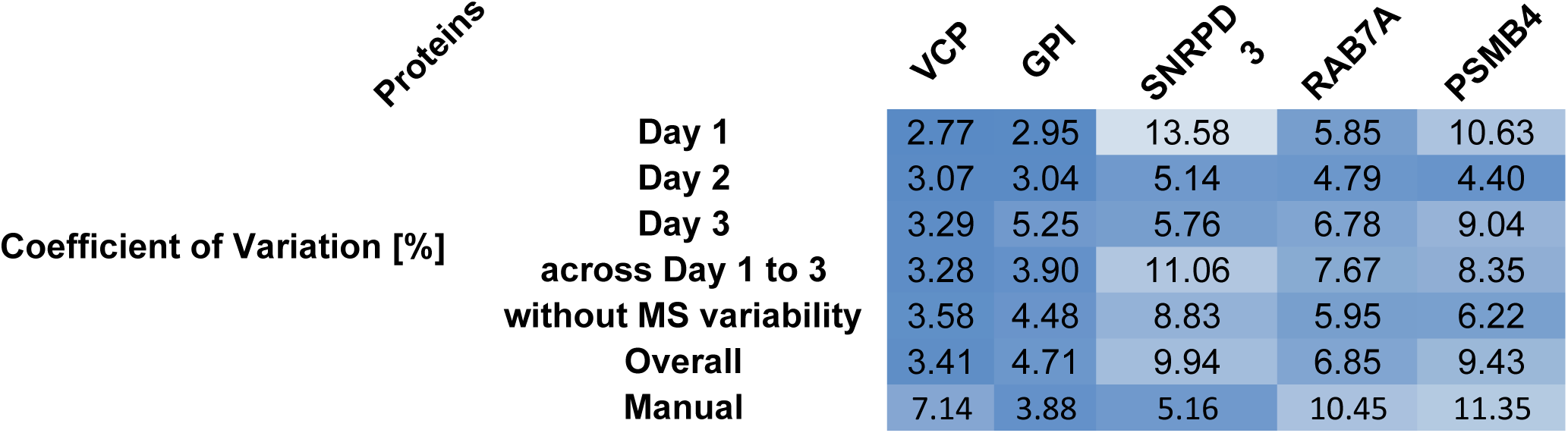

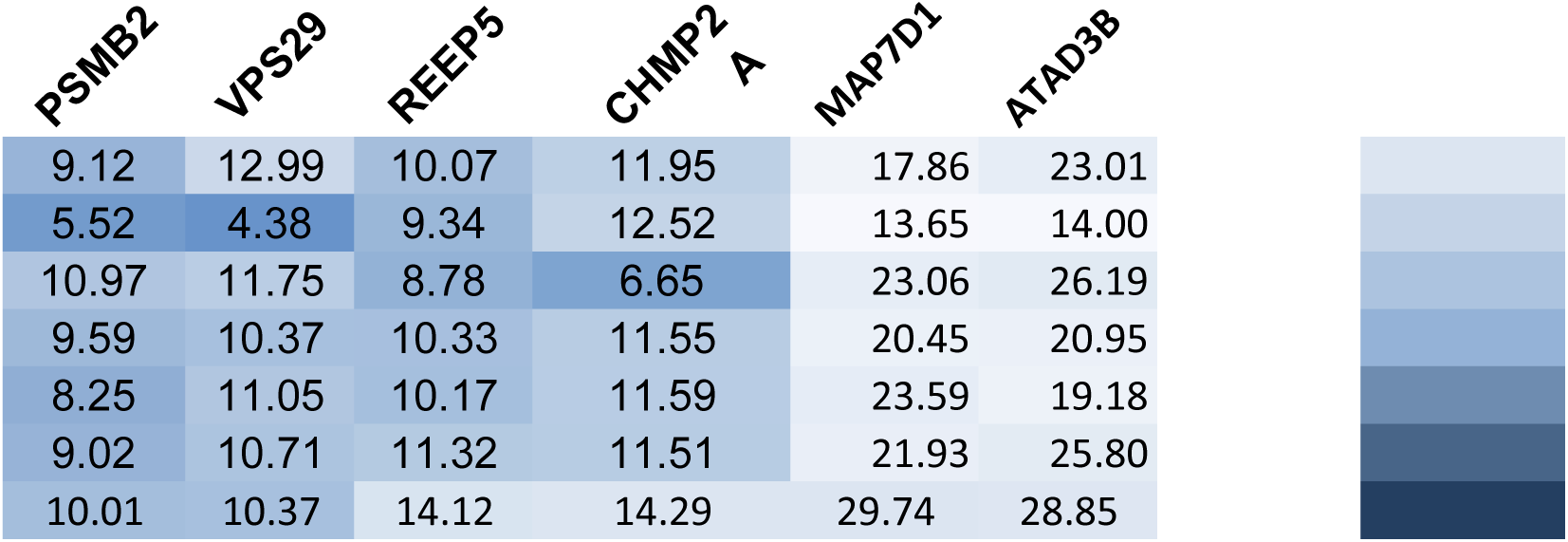

## Supplementary Protocols

***Supplementary Protocol A. Single-step reduction and alkylation with TCEP/ CAA plus core autoSP3 clean-up***. Detailed description of main protocol tasks with step-by-step liquid handling, plate movements, and process time.

***Supplementary Protocol B. Two-step reduction and alkylation with e.g. DTT/ CAA plus core autoSP3 clean-up***. Detailed description of main protocol tasks with step-by-step liquid handling, plate movements, and process time.

***Supplementary Protocol C. Core autoSP3 clean-up (omitting automated reduction and alkylation)***. Detailed description of main protocol tasks with step-by-step liquid handling, plate movements, and process time.

***Supplementary Protocol D. Acidification and recovery of peptides to a new sample plate***. Detailed description of main protocol tasks with step-by-step liquid handling, plate movements, and process time.

